# mRNA 3′ UTRs direct microRNA degradation to participate in imprinted gene networks and regulate growth

**DOI:** 10.1101/2025.11.06.686990

**Authors:** Daniel H. Lin, Lara E. Elcavage, Ekaterina Khalizeva, David P. Bartel

## Abstract

MicroRNAs direct downregulation of target mRNAs. Sometimes, however, this regulatory paradigm inverts, and a target RNA triggers the degradation of a microRNA. This target-directed microRNA degradation (TDMD) requires ZSWIM8. *Zswim8*^−/−^ mice exhibit reduced growth and perinatal lethality, accompanied by stabilization of dozens of microRNAs. Nonetheless, studies of TDMD function in mammals have been limited because only two TDMD-triggering RNAs have been identified in mice. Here, we computationally identify and validate five new TDMD-triggering sites in mouse models. One site in *Atp6v1g1* and two in *Lpar4* direct degradation of miR-335-3p, which shows that in mammals, two sites in the same transcript, and multiple sites in different transcripts, can collaborate to destabilize a microRNA. Moreover, sites in *Plagl1* and *Lrrc58* direct degradation of miR-322 and miR-503, respectively. Mice lacking the *Plagl1* and *Lrrc58* sites exhibit reduced growth, demonstrating that target-directed degradation of miR-503 and miR-322 promotes mammalian growth. Both miR-335-3p and *Plagl1* are maternally imprinted, implying that they participate in parental conflict, but their corresponding triggers or target microRNA partner are not imprinted. Thus, 3′ UTRs directly participate in parental conflict by engaging TDMD to access an additional layer of regulation within a network of imprinted and biallelic genes.

## Introduction

MicroRNAs (miRNAs) are ∼22-nt RNAs that specify post-transcriptional gene repression (Bartel 2018). Mammalian genomes encode >500 miRNA genes (Fromm et al. 2015, 2020, 2022; Clarke et al. 2025), and each miRNA can target hundreds of different mRNAs across the transcriptome, such that in aggregate, most human mRNAs are conserved targets of at least one miRNA (Friedman et al. 2009). Reinforcing this broad role for miRNAs, mice that lack either individual miRNAs or several members of the same miRNA family possess a wide spectrum of phenotypes (Bartel 2018). Precise control of miRNA expression is also critical across development, as overexpression of miRNAs can also result in developmental defects or disease (Kloosterman and Plasterk 2006; Sayed and Abdellatif 2011; Concepcion et al. 2012).

Each miRNA acts in concert with an Argonaute (AGO) protein, which envelops the miRNA and shapes its interactions with target mRNAs (Schirle and MacRae 2012; Elkayam et al. 2012; Faehnle et al. 2013; Schirle et al. 2014). The miRNA 5′ and 3′ ends each bind specific pockets in AGO, which largely shields these ends from cellular exonucleases. As a result of this protection, miRNAs are typically very stable, with median half-lives of 34 h in mouse embryonic fibroblasts (MEFs), compared to median half-lives of 2 h for mRNAs (Kingston and Bartel 2019; Eisen et al. 2020). Within the Ago– miRNA complex, the miRNA recognizes mRNA targets, primarily through base pairing between nucleotides 2–7 of the miRNA (called the miRNA seed) and sites within the 3′ UTRs of the mRNAs.

This seed pairing can be augmented on either end by additional pairing to miRNA nucleotide 8 or the presence of a target A nucleotide across from miRNA nucleotide 1 (Lewis et al. 2005; Grimson et al. 2007; Schirle et al. 2014, 2015). Although not typically observed, additional pairing between the target and the 3′ region of the miRNA can supplement seed pairing to increase affinity for the target (supplementary pairing) or compensate for defects in base pairing in the seed region (compensatory pairing) (Brennecke et al. 2005; Grimson et al. 2007; Bartel 2009). Stable association of the Ago– miRNA complex with the target RNA typically leads to recruitment of TNRC6, which in turn recruits deadenylation complexes that shorten the mRNA poly(A) tail. Pairing to the central nucleotides 9 and 10 is usually sterically occluded by AGO, but in conjunction with a conformational change, complete complementarity between the miRNA and the target activates AGO2 endonuclease activity, which slices the target RNA (Hutvágner and Zamore 2002; Sheu-Gruttadauria et al. 2019; Mohamed et al. 2025).

In rare cases, an inversion of this regulatory paradigm can occur, wherein the target RNA triggers selective decay of the miRNA. This target-directed miRNA degradation (TDMD) is mediated by a Cullin-Ring E3 ubiquitin ligase complex constituted by the substrate receptor ZSWIM8, adaptor proteins ELOB and ELOC, and CUL3, which recruits additional ubiquitylation factors (Han et al. 2020; Shi et al. 2020). Polyubiquitylation of AGO causes its degradation by the proteasome, leaving the miRNA susceptible to cellular nucleases while freeing the TDMD-triggering RNA (or TDMD trigger) to direct additional cycles of TDMD (Han et al. 2020; Shi et al. 2020).

TDMD was first observed in response to either synthetic RNA targets with extensive complementarity (Ameres et al. 2010) or viral transcripts that direct the degradation of cellular miRNAs that would otherwise slow viral replication (Cazalla et al. 2010; Libri et al. 2012; Marcinowski et al. 2012; Lee et al. 2013). More recently, TDMD has been described for a handful of endogenously encoded sites — e.g., two in mice, seven in Drosophila, and one in worms (Bitetti et al. 2018; Kleaveland et al. 2018; Kingston et al. 2022; Sheng et al. 2023; Grimme et al. 2025; Hiers et al. 2025)— but molecular analyses of the effects of ZSWIM8 disruption in diverse contexts suggest the existence of many more that have yet to be found (Han et al. 2020; Shi et al. 2020, 2023; Kingston et al. 2022; Jones et al. 2023; Stubna et al. 2025).

In mice, loss of ZSWIM8 results in heart and lung development defects, reduced growth, and perinatal lethality, suggesting a critical role for TDMD in development (Jones et al. 2023; Shi et al. 2023). Accompanying these phenotypes, >40 miRNAs increase in abundance in various tissues of the mouse embryo, suggesting they may be substrates of endogenous TDMD (Jones et al. 2023; Shi et al. 2023). However, only two murine transcripts have been identified and validated as endogenous transcripts that specify TDMD in vivo: the *Nrep* mRNA directs miR-29 degradation, and the Cyrano long noncoding RNA (lncRNA) directs miR-7 degradation (Bitetti et al. 2018; Kleaveland et al. 2018). *Serpine1* and *BCL2L11* mRNAs have also been reported as TDMD triggers for miR-30c/30e and miR-221/222, respectively, but these miRNAs are not significantly upregulated in *Zswim8^-/-^* embryos (Ghini et al. 2018; Li et al. 2021; Jones et al. 2023; Shi et al. 2023). Determining whether the remaining ZSWIM8-sensitive miRNAs are also TDMD substrates requires identification of transcripts that direct their degradation.

The most notable feature of the *Cyrano* and *Nrep* sites that trigger TDMD is their extensive complementarity to the 3′ region of their respective miRNAs (Bitetti et al. 2018; Kleaveland et al. 2018). Indeed, these sites are the most extensively paired sites in the transcriptome for either miRNA (Figure S1A–B). The prevailing paradigm is that this extensive, TDMD-triggering 3′ complementarity is distinct from 3′ supplementary pairing and induces a conformational change in the Ago–miRNA– target ternary complex that can be specifically recognized by ZSWIM8, but the molecular details of such recognition are poorly understood (Sheu-Gruttadauria et al. 2019; Buhagiar and Kleaveland 2024). In support of this paradigm, mutations that attenuate either seed pairing or extensive complementarity disrupt TDMD (Kleaveland et al. 2018; Sheu-Gruttadauria et al. 2019).

Although most genes are expressed equally from both sets of chromosomes, a few are preferentially expressed from one allele, which results in an asymmetric contribution from the paternal and maternal alleles (Surani et al. 1984; McGrath and Solter 1984; Cattanach and Kirk 1985; Peters 2014; Cleaton et al. 2014; Tucci et al. 2019). These imprinted genes gain robust epigenetic modifications in either the maternal or paternal germline that result in their silencing in the offspring. In mice, more than 200 such imprinted genes have been reported (Tucci et al. 2019). Among vertebrates, the process of imprinting is restricted to therian mammals, suggesting a link between imprinting and resource allocation in utero (Cleaton et al. 2014; Tucci et al. 2019). Kinship theory posits that the interests of the two parental genomes within offspring can conflict with respect to the use of maternal resources to support the growth of the fetus or newborn (Haig and Westoby 1989; Moore and Haig 1991; Tucci et al. 2019). For example, a murine gene that causes more maternal resources to be used to promote growth of the fetus or pup at the expense of either littermates or future litters will favor the interests of the paternal genome, whereas a gene that saves maternal resources for the benefit of littermates or future litters will favor the interests of the maternal genome. In this scenario in which the fitness of an allele can differ depending on its inheritance pattern, imprinting provides the means by which the parents can favor the interests of their chromosomes (Peters 2014; Tucci et al. 2019). In support of this hypothesis, many maternally imprinted, paternally expressed genes increase the allocation of maternal resources to promote growth of the fetus or newborn, whereas paternally imprinted, maternally expressed genes limit growth (Peters 2014).

Interestingly, miRNAs that emerged around the time of the last common ancestor of placental mammals are highly enriched for imprinted loci, implying that miRNAs were enlisted early to do battle in this conflict. Indeed, of the 88 miRNA families conserved among placental mammals but absent in fish, more than 40% are imprinted, and of these, at least 40% are ZSWIM8-sensitive (Jones et al. 2023; Shi et al. 2023). TDMD may also have been co-opted into this genomic conflict.

Here, we identified four additional endogenous triggers for three ZSWIM8-sensitive miRNAs, thereby tripling the number of known triggers in mammals. Of these four triggers, one is maternally imprinted and two direct the destruction of a maternally imprinted miRNA, thereby strengthening the link between the TDMD pathway and genomic imprinting.

## Results

### Computational identification of triggers for ZSWIM8-sensitive miRNAs

Although each miRNA typically possesses hundreds to thousands of potential target sites throughout the transcriptome, few of these sites, if any, trigger TDMD, creating a fundamental challenge as to how to identify the exceedingly rare sites that are effective TDMD triggers. The first endogenous TDMD triggers were identified through their canonical seed pairing combined with their unusually extensive pairing to the miRNA 3′ region. Subsequent studies have focused on sites that resemble these few known and characterized examples. These sites have a few unifying features: (1) canonical seed pairing, (2) extensive complementarity to the miRNA 3′ region, (3) high gene expression, and (4) evolutionary conservation.

Searching for sites with these characteristics has been most productive in Drosophila, identifying triggers for about half of the Dora-sensitive miRNAs in Drosophila S2 cells (Kingston et al. 2022). Accordingly, we used a similar scheme to search for candidate TDMD sites within the transcriptome of MEFs. For one branch of this search, we considered only sites with conserved miRNA seed matches in mRNA 3′ UTRs (Figure S1A). In another branch we considered all sites with canonical seed matches in mRNA 3′ UTRs and annotated lncRNAs (Figure S1B). Sites were filtered for expression in MEFs, and extensively complementary targets were prioritized with a scoring scheme that (1) awarded points for Watson–Crick–Franklin base pairing to the 3′ region of the miRNA, (2) penalized non-contiguous base pairing, (3) awarded additional points for pairing to nucleotides near the end of the miRNA, and (4) penalized large bulges or internal loops in predicted pairing between the miRNA and target. As an orthogonal metric for pairing quality, we also predicted the folding energy for pairing between the miRNA-binding site and the miRNA 3′ region, as was done in Drosophila (Kingston et al. 2022). Both approaches confirmed that the previously characterized miRNA-binding sites in *Cyrano* and *Nrep* possessed exceptional pairing to the 3′ region of miR-7 and miR-29b, respectively (Figure S1A–B). Top-scoring candidates for other miRNAs were tested by knocking down or knocking out the site predicted to direct miRNA degradation and monitoring the effect on the miRNA. This approach identified triggers for miR-335-3p, miR-322, and miR-503.

### Multiple TDMD sites, from the same and different trigger transcripts, direct degradation of miR-335-3p

Mouse miR-335-3p is one of the most potently and broadly upregulated miRNAs in *Zswim8*^-/-^ embryonic tissues (Jones et al. 2023; Shi et al. 2023). The *Mir335* pri-miRNA hairpin is located within an intron of the maternally imprinted gene *Mest* and is therefore only expressed from the paternal allele (Figure 1A) (Hiramuki et al. 2015). Typically, a miRNA is generated from an RNA hairpin that is processed in successive cleavage steps to yield a duplex of two ∼22-nt strands derived from the 5′ and 3′ arms of the hairpin (Bartel 2018; Kim et al. 2025). One strand of each duplex associates with an AGO protein and serves as the functional guide RNA, whereas the other strand, known as the passenger strand, is discarded and rapidly degraded. Most miRNA duplexes exhibit a bias as to which strand associates with AGO and which one is discarded, and the relative accumulation of the guide and passenger strands reflects this bias (Bartel 2018). For miR-335, TDMD also helps to determine the more prominent strand (Jones et al. 2023; Shi et al. 2023). For this miRNA, the strand from the 3′ arm of the hairpin, designated miR-335-3p, typically accumulates at much lower levels than its co-produced strand from the 5′ arm of the hairpin, miR-335-5p, but this ratio is a consequence of the potent, strand-specific turnover of miR-335-3p by TDMD (∼7.4-fold in MEFs; up to ∼17.8-fold in embryonic day 18.5, E18.5, stomach) (Figure 1A–B) (Shi et al. 2023). Thus, miR-335-3p is the more abundant strand in *Zswim8* knockout tissues — a rare example in which TDMD causes apparent “arm switching”.

**Figure 1.**
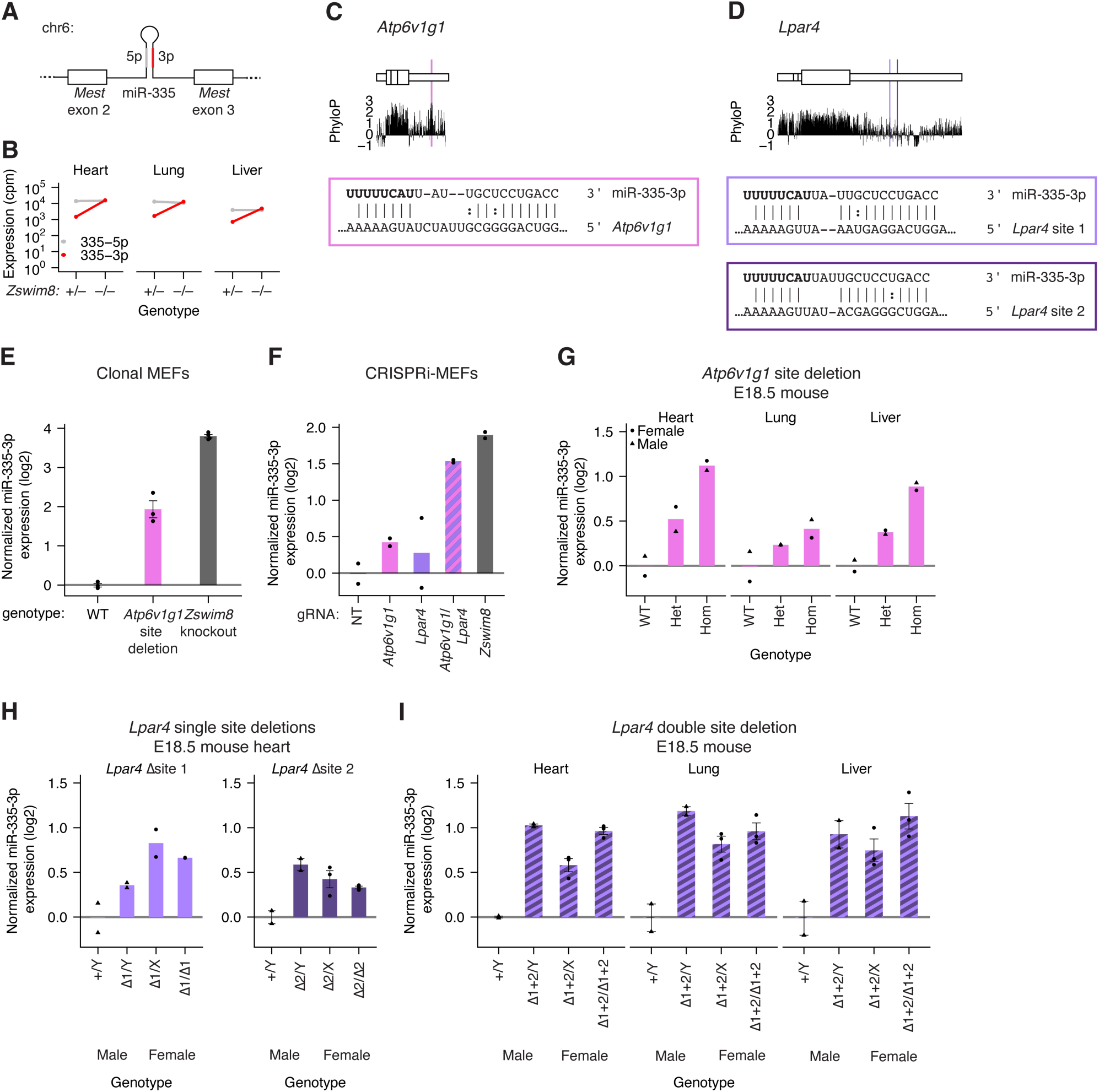
Sites in *Atp6v1g1* and *Lpar4* 3′ UTRs collaborate to mediate TDMD in MEFs and in vivo. A) Structure of the *Mest* gene, which harbors the *Mir335* gene within intron 2. B) Expression of miR-335-3p and miR-335-5p strands in embryonic day 18.5 (E18.5) heart, lung, and liver of *Zswim8*^-/+^ and *Zswim8^-^*^/-^ (cpm, counts per million miRNA reads) (Shi et al. 2023). C) mRNA context, evolutionary conservation, and base-pairing diagram for the miR-335-3p trigger site in *Atp6v1g1. Top*: The *Atp6v1g1* mRNA, depicting coding sequence in larger rectangles and UTRs in smaller rectangles. Vertical lines indicate exon boundaries. *Middle*: PhyloP score from a mammalian 60-way alignment (Yang 1995; Felsenstein and Churchill 1996; Murphy et al. 2001; Chiaromonte et al. 2002; Schwartz et al. 2003; Kent et al. 2003; Blanchette et al. 2004; Siepel and Haussler 2005; Pollard et al. 2010) plotted in 5-nt bins. *Bottom*: Pairing diagram depicting complementarity between miR-335-3p and its trigger site in the *Atp6v1g1* 3′ UTR. Vertical lines indicate W–C–F pairing; colon indicates G:U wobble pairing. D) mRNA context, evolutionary conservation, and base-pairing diagrams for the miR-335-3p trigger sites in *Lpar4*; otherwise as in C. E) Function of the miR-335-3p trigger site in *Atp6v1g1*. Plotted is quantification of miR-335-3p, as measured by small RNA sequencing (sRNA-seq), from clonal MEF cell lines with either homozygous deletion of the miR-335-3p trigger site or *Zswim8* knockout. To account for clone-to-clone variability in miR-335 production, the depth-normalized expression of miR-335-3p was normalized to the expression of its co-transcribed miR-335-5p strand. Each point represents the fold-change of normalized expression relative to the mean of normalized expression in WT samples (*n* = 3–4 clonal lines per genotype; error bars, standard error). F) Function of the *Atp6v1g1* and *Lpar4* trigger transcripts. Plotted is quantification of miR-335-3p expression, as measured by sRNA-seq following CRISPRi knockdown of either *Atp6v1g1*, *Lpar4*, both mRNAs, or *Zswim8* (*n* = 2 biological replicates). Otherwise, this panel is as in E. G) In vivo TDMD activity of the miR-335-3p trigger site in *Atp6v1g1.* Shown is quantification by northern blot of miR-335-3p expression, normalized to miR-335-5p expression, in E18.5 heart, lung, and liver of mice harboring deletions of the miR-335-3p TDMD site in *Atp6v1g1* (*Atp6v1g1*^-50^). Circles represent female animals; triangles represent male animals (*n* = 2 replicates per tissue for each genotype). H) In vivo TDMD activity of site 1 or site 2 in *Lpar4* (site 1 mutant: *Lpar4*^-173^; site 2 mutant: Lpar4^-69+36^; *n* = 2–3 replicates per tissue for each genotype and sex). Otherwise, this panel is as in G. H) In vivo activity of both miR-335-3p trigger sites in *Lpar4* (site 1 and 2 mutant: *Lpar4*^-197^; *n* = 2 replicates per tissue for male samples and *n* = 3 replicates per tissue for female samples). Otherwise, this panel is as in G.

We identified a site in the 3′ UTR of *Atp6v1g1* and two sites in *Lpar4* as the top-scoring candidate TDMD sites for miR-335-3p (Figure S1A–B; Figure 1C–D). The potential role of the *Atp6v1g1* site in MEFs was tested by Cas9-mediated deletion of the site using two flanking guide RNAs, which yielded ∼50-nt deletions encompassing the miR-335-3p site. Although substantial clone-to-clone variability in expression was observed for both strands of miR-335, presumably caused by differences in production of the miRNA hairpin, the post-transcriptional effects of *Atp6v1g1* site deletion on miR-335-3p could be isolated by normalizing to the expression of the co-produced miR-335-5p strand. After this normalization, the effect of *Atp6v1g1* site deletion on miR-335-3p expression was only ∼25% of the effect of *Zswim8* knockout, suggesting that another trigger was acting partially redundantly in MEFs (Figure 1E, S2A). Suspecting that the sites in *Lpar4* might be responsible for the remaining effect, a similar deletion strategy was attempted for these two sites, but we were unable to obtain clones in which the *Lpar4* sites were deleted on all alleles. Therefore, we turned to CRISPRi knockdown experiments. In these experiments, we observed weak effects when either *Atp6v1g1* or *Lpar4* were individually targeted but nearly the full effect of the *Zswim8* knockdown when both genes were targeted (Figure 1F, S2B). We conclude that these two transcripts collaborate to direct the degradation of miR-335-3p in MEFs.

Although *Atp6v1g1* and *Lpar4* have similar contributions to miR-335-3p turnover, they are expressed at substantially different levels. In MEFs, *Atp6v1g1* expression is ∼42-fold greater than *Lpar4* expression, as measured by RNA-seq (179 vs 4.2 transcripts per million (TPM)) (Figure S2C). Even after accounting for the presence of two sites per *Lpar4* transcript, this difference in expression levels suggests more efficient turnover of miR-335-3p by *Lpar4* than by *Atp6v1g1*.

To more thoroughly validate the three TDMD sites in *Atp6v1g1* and *Lpar4*, we generated *Atp6v1g1* and *Lpar4* mutant mice with targeted deletions of the TDMD sites. The *Atp6v1g1*^-50^ mice possessed a 50-nt deletion encompassing the miR-335-3p site (Figure S3A). The allelic series of *Lpar4* mutant mice possessed deletions that disrupted either an individual site (site 1 mutant: *Lpar4*^-173^; site 2 mutant: Lpar4^-69+36^) or both sites (*Lpar4*^-197^) (Figure S3B).

We bred mice heterozygous for the *Atp6v1g1*^-50^ allele and examined miR-335-3p levels at embryonic day 18.5 (E18.5). Homozygous deletion of the TDMD site in *Atp6v1g1* resulted in an increase in miR-335-3p levels in all three embryonic tissues examined — heart, lung, and liver (Figure 1G, S3C). Heterozygous deletion of the site resulted in an intermediate level of miR-335-3p elevation, indicating a dosage-dependent effect of TDMD activity by *Atp6v1g1*. The increases in miR-335-3p observed in homozygous *Atp6v1g1*^-50^ tissues represented 15%, 6%, and 16% of that reported in *Zswim8^-/-^*tissues compared to *Zswim8^-/+^*tissues (Heart 2.4-fold vs. 10.5-fold; Lung: 1.4-fold vs. 7.8-fold; Liver: 1.9-fold vs. 6.6-fold) (Shi et al. 2023), which suggested that at least one additional trigger — perhaps *Lpar4* — might be required to achieve the full effect of ZSWIM8.

To assess the contribution of TDMD sites in *Lpar4* to miR-335-3p degradation, we bred hemizygous males with heterozygous females for each of the alleles in the allelic series. Deletion of either individual TDMD site in *Lpar4* caused 1.3-fold and 1.6-fold increases in miR-335-3p levels in male E18.5 heart for site 1 and site 2, respectively (Figure 1H, S3D–E). Deletion of both TDMD sites in *Lpar4* caused a greater increase in miR-335-3p levels (2.0-fold in heart, 2.3-fold in lung, 1.7-fold in liver), accounting for 11–19% of the reported effect in *Zswim8^-/-^* tissues (Shi et al. 2023) (Figure 1I, S3F).

These observations in *Atp6v1g1* and *Lpar4* mutant mice confirmed the independent function of these two triggers for miR-335-3p TDMD in vivo. Consistent with observations in cell culture, these two transcripts contributed similarly to the degradation of miR-335-3p in animals, despite vast differences in their expression (Shi et al. 2023) (Figure S2C), suggesting the existence of unknown factors that might enhance the potency of some triggers.

Overall, these results showed that a miRNA can be targeted by multiple partially redundant TDMD sites — both in a single transcript or across multiple transcripts. In the case of the exceptionally sensitive miRNA miR-335-3p, we could detect partial effects for perturbation of individual TDMD sites. However, in the more typical scenario, in which the miRNA is more modestly ZSWIM8-sensitive, the effect of a single site would presumably be more difficult to detect, and simultaneous perturbation of multiple candidate sites might be required to reveal their activity.

### *Plagl1* and *Lrrc58* direct degradation of miR-322 and miR-503

We also considered the potential TDMD triggers of miR-322 and miR-503, two miRNAs that have unusually short half lives in mammalian cells (Rissland et al. 2011; Kingston and Bartel 2019). These two miRNAs are members of the extended miR-15/16 seed family and are co-expressed from a miRNA cluster located on the X chromosome (Figure 2A). miR-322 has the same sequence in its seed region as other members of the miR-15/16 seed family, whereas miR-503 differs at position 8 of the seed region. As a result, these miRNAs are predicted to target partially overlapping but distinct cohorts of target mRNAs (Agarwal et al. 2015; McGeary et al. 2019). Despite these similarities in the seed region, the sequences of these miRNAs diverge substantially in their 3′ regions, implying that different trigger sites might direct their degradation. Indeed, a computational search for extensively paired conserved sites for miR-322 and miR-503 identified sites in the 3′ UTRs of different mRNAs — with a miR-322 site in the *Plagl1* 3′ UTR and a miR-503 site in the *Lrrc58* 3′ UTR found among the top hits (Figure 2B–C, S1A–B). Notably, the proposal that pairing to the site in *Lrrc58* directs degradation of miR-503 is fully consistent with results of scanning mutagenesis that identify the nucleotides of miR-503 responsible for its destabilization (Rissland et al. 2011).

**Figure 2.**
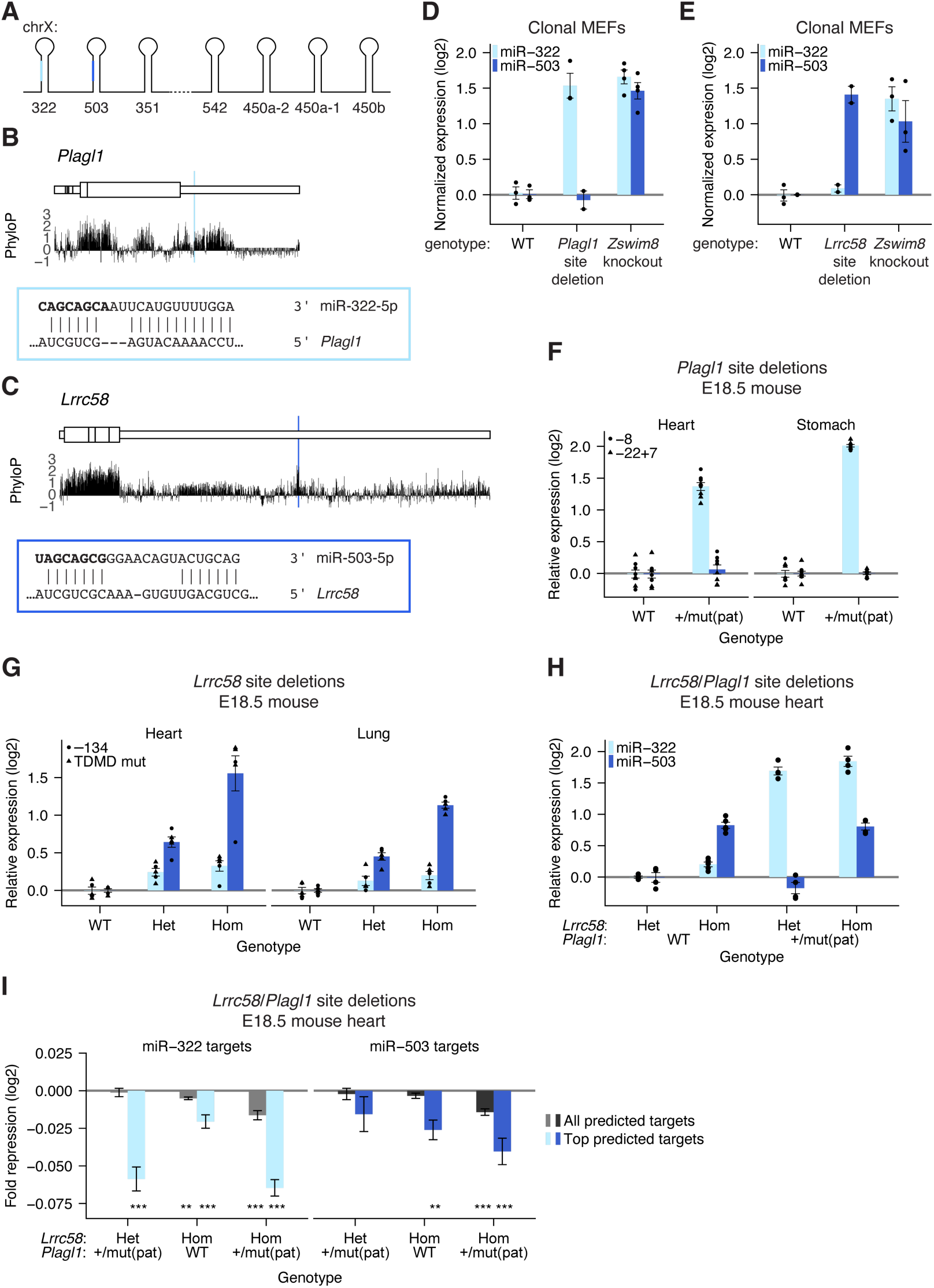
Sites in the 3′ UTRs of *Plagl1* and *Lrrc58* are the major TDMD triggers of miR-322-5p and miR-503-5p, respectively. A) Organization of the X-linked cluster of seven miRNA genes, including genes for miR-322 and miR-503. The dashed line between the *Mir351* and *Mir542* hairpins denotes a longer genomic distance and loss of coordinated expression. B) mRNA context, evolutionary conservation, and base-pairing diagram for the miR-322 trigger site within *Plagl1*; otherwise, as in Figure 1C. C) mRNA context, evolutionary conservation, and base-pairing diagram for the miR-503 trigger site within *Lrrc58*; otherwise, as in Figure 1C. D) Function of the miR-322 trigger site in *Plagl1*. To account for clone-to-clone variability in cluster expression, the depth-normalized expression of miR-322-5p was first normalized to the expression of co-transcribed miRNAs miR-322-3p, miR-503-3p, and miR-351-3p; (*n* = 3–4 clonal lines per genotype); otherwise, as in Figure 1E. E) Function of the miR-503 trigger site in *Lrrc58* (*n* = 2–3 clonal lines per genotype); otherwise, as in D. F) In vivo activity of the miR-322 trigger site in *Plagl1*. Shown is quantification by northern blot of miR-322-5p or miR-503-5p expression in E18.5 heart and lung tissue of *Plagl1*-mutant mice harboring an 8-nt deletion or 22-nt deletion/7-nt insertion within the miR-322 TDMD site (*n* = 8 replicates per tissue for each genotype). G) In vivo activity of the miR-503 trigger site in *Lrrc58*. Quantification of miR-322-5p or miR-503-5p expression by northern blot in E18.5 heart and lung tissue of *Lrrc58* mutant mice harboring either a 134-nt deletion or a precise mutation of the miR-503 TDMD trigger site (*n* = 5 replicates per tissue for each genotype). H) In vivo activity of both the miR-322 trigger site in *Plagl1* and the miR-503 trigger site in *Lrrc58*. Quantification of miR-322-5p or miR-503-5p expression by northern blot in E18.5 heart tissue of mutant mice harboring deletions of the miR-322 and miR-503 TDMD trigger sites in *Plagl1* and *Lrrc58*. Genotypes are indicated below the x-axis (*Lrrc58* het, heterozygous mutant *Lrrc58*; *Lrrc58* hom, homozygous mutant *Lrrc58*; *Plagl1* WT, wild-type *Plagl1*; *Plagl1* +/mut(pat), heterozygous mutant *Plagl1* with paternal inheritance; *n* = 4–5 replicates for each genotype.) I) The influence of trigger sites within *Plagl1* and *Lrrc58* on levels of predicted miR-322-5p or miR-503-5p targets in vivo. Two sets of targets were analyzed: all predicted targets and top predicted targets (top 10% of predicted targets, as determined by TargetScan) (Agarwal et al. 2015). For each set of predicted targets, a set of nontarget transcripts, matched for 3′ UTR length, was sampled at a 1:1 ratio, and the distributions of log2 fold changes of mutant mice compared to *Plagl1* WT, *Lrrc58* het littermates (genotypes indicated along the x-axis, as in H) were compared between predicted target and nontarget cohorts. The repression metric plotted is the difference between median log2 fold change (mutant/WT) of the target cohort and that of the nontarget cohort. This analysis was repeated 20 additional times, sampling new nontarget cohorts with each iteration. Plotted is the mean repression metric and median *P* value across the 21 iterations (error bars, standard deviation; * *P* <0.05, ** *P* <0.005, *** *P* <0.0005, Mann–Whitney U test comparing the distribution for predicted targets and that of their nontarget cohort).

Disruption of the site in *Plagl1* caused miR-322 to accumulate to levels comparable to *Zswim8* knockout, after accounting for clone-to-clone variability in cluster expression (Figure 2D, S4A). Likewise, disruption of the site in *Lrrc58* caused miR-503 to accumulate to levels comparable to *Zswim8* knockout, again after accounting for clone-to-clone variability in cluster expression (Figure 2E, S4B). Thus, in MEFs, these two mRNAs were the major TDMD triggers for their respective miRNAs.

We next assessed the extent to which *Plagl1* and *Lrrc58* triggered TDMD in the mouse embryo. We generated two *Plagl1* lines with either an 8-nt deletion (*Plagl1*^-8^) or a 22-nt deletion/7-nt insertion (*Plagl1*^-22+7^) that disrupted complementarity to the 3′ region of miR-322 (Figure S4C). Because *Plagl1* is maternally imprinted, heterozygous mice with a paternally inherited mutant allele (*Plagl1*^+/mut(pat)^) express only the mutant allele (Kamiya 2000; Piras et al. 2000). The increased miR-322 accumulation in *Plagl1*^+/mut(pat)^ embryonic tissues approached that observed when comparing *Zswim8*^-/-^ embryonic tissues with *Zswim8*^-/+^ embryonic tissues (Heart: 2.6-fold vs. 2.7-fold; Stomach: 4.0-fold vs. 5.7-fold) (Shi et al. 2023) (Figure 2F, S4D–E).

Similarly, for *Lrrc58*, we generated three mouse lines containing either four substitutions in the miR-503 TDMD site (*Lrrc58*^TDMD^ ^mut^), or 134- or 402-nt deletions encompassing the site (*Lrrc58*^-134^ and *Lrrc58*^-402^, respectively) (Figure S4F). Disruption of the site via targeted substitutions or deletion of the entire site resulted in similar upregulation of miR-503 in E18.5 heart and lung, which resembled or surpassed that reported for the increased accumulation in *Zswim8^-/-^* tissues compared to *Zswim8^-/+^* tissues (Heart: 3.5-fold vs. 2.5-fold; Lung 3.6-fold vs. 1.8-fold) (Shi et al. 2023) (Figure 2G, S4G–H). Similar to observations in *Atp6v1g1*^-50^ embryonic tissue, miR-503 was elevated by an intermediate amount in mice heterozygous for the *Lrrc58* site mutation, demonstrating dosage-dependent TDMD by *Lrrc58* (Figure 2G).

The miR-322 fold change in *Plagl1* mutant tissues was somewhat less than that observed when comparing *Zswim8*^-/-^ with *Zswim8*^-/+^ tissues (Shi et al. 2020). Perhaps explaining this result, miR-322 is weakly but consistently upregulated in *Lrrc58* mutant samples (Figure 2G), and miR-322 accumulation increased 3.6-fold in *Plagl1*^+/–22+7(pat)^, *Lrrc58*^-134/–134^ double-mutant heart samples (Figure 2H, S5A). Thus, whereas *Plagl1* and *Lrrc58* were the dominant TDMD triggers for miR-322 and miR-503, respectively, in these embryonic tissues, *Lrrc58* appeared to have weak TDMD activity for miR-322 as well.

Consistent with upregulation of miR-322 and miR-503 in the TDMD site mutants, predicted targets of miR-322 and miR-503 were more repressed in mutant E18.5 heart tissue (Figure 2I, S5B–C). For each miRNA, the greatest repression of predicted targets was observed in *Plagl1*/*Lrrc58* double mutants, which was consistent with the partially overlapping targetome of these two related miRNAs.

### ZSWIM8 sensitivity is often, but not always, conserved among mammals

Although more than 50 miRNAs increase in expression in murine *Zswim8^-/-^* embryonic tissues or cell lines, many fewer miRNAs have been observed to increase in expression in human cell lines — a total of 12 miRNAs were confidently annotated as increasing across five cell lines (Shi et al. 2020). The simplest explanation for this difference is that the human analyses have been performed in more restricted contexts — primarily in cancer cell lines. If so, perturbation of *ZSWIM8* in human fibroblast cell lines, which more closely resemble the previously assessed mouse fibroblast cell lines, might identify more TDMD substrates in human cells. Indeed, sequencing miRNAs from *ZSWIM8* knockout HFF-1, IMR90, and BJ human fibroblast lines identified 27, 34, and 34 upregulated miRNAs, respectively, after stringent statistical filtering (Figure 3A–C, S6A–C; Table S1) (Wang and Bartel 2023). Altogether, we identified 47 unique ZSWIM8-sensitive miRNAs, expanding the cohort of miRNAs reported to be ZSWIM8-sensitive in human cells by 39 (Figure 3D–E). Of the 41 mouse ZSWIM8-sensitive miRNAs with clear human homologs, 18 miRNAs passed statistical thresholds in at least one human fibroblast line (Figure 3F, Table S2). Moreover, some that did not pass the statistical thresholds were nonetheless mildly upregulated (Figure S6D–F). Thus, we expect that this analysis provided a conservative lower bound on conservation of TDMD between the two species.

**Figure 3.**
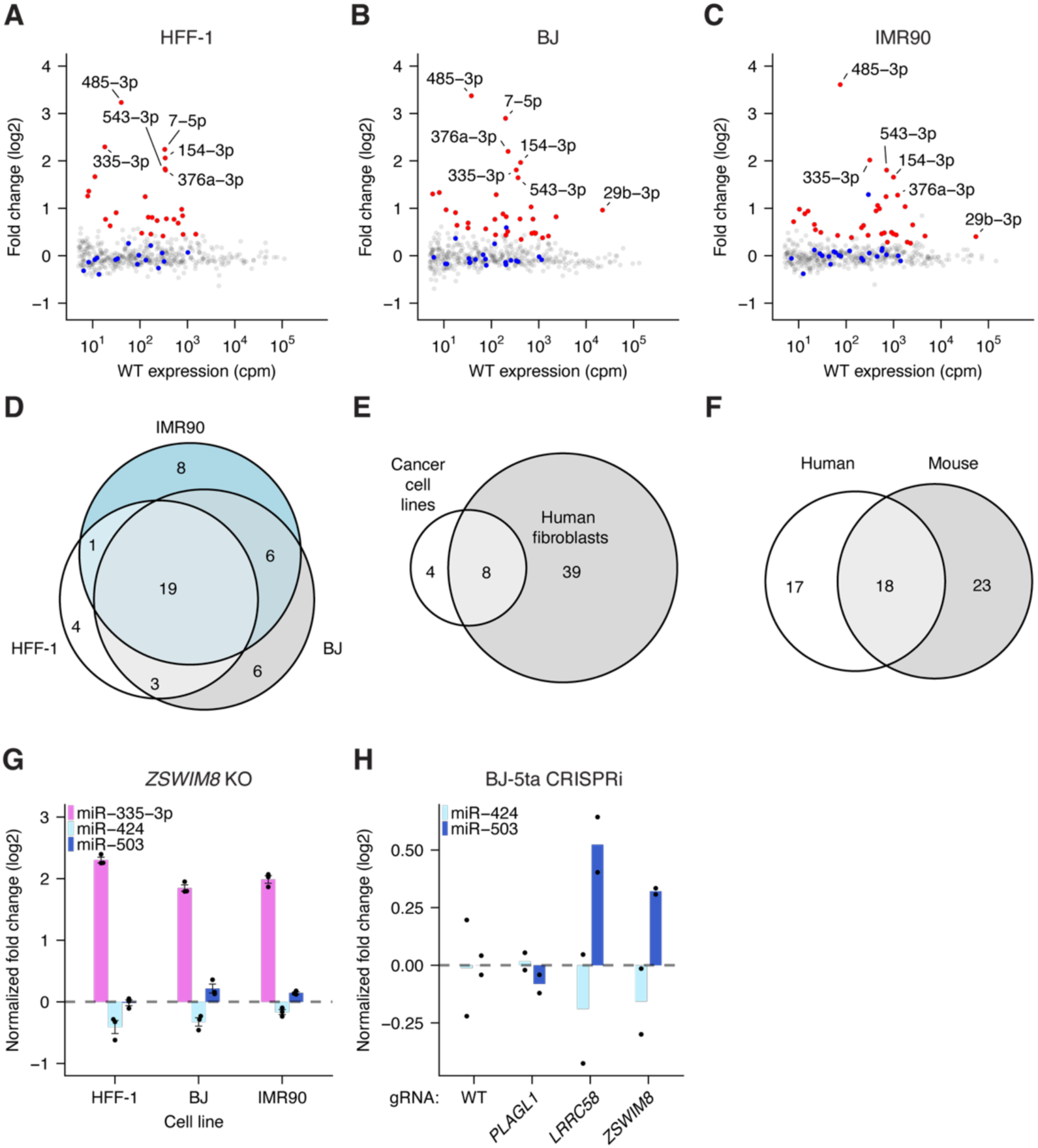
Evolutionary conservation of ZSWIM8 sensitivity. A) ZSWIM8 sensitivity in HFF-1 cells. Plotted are fold changes of miRNA levels upon polyclonal knockout of *ZSWIM8*, as measured by sRNA-seq with analysis by DESeq2 (*n* = 3 biological replicates) (Love et al. 2014). In red are values for which ZSWIM8 sensitivity was statistically significant (Wang and Bartel 2023) (Table S1). B) ZSWIM8 sensitivity in BJ cells; otherwise, as in A. C) ZSWIM8-sensitivity in IMR90 cells; otherwise, as in A. D) Overlap of ZSWIM8-sensitive miRNAs in three human fibroblast cell lines. E) Overlap of miRNAs found to be ZSWIM8-sensitive in human fibroblasts with those found to be sensitive in human cancer cell lines (Shi et al. 2020). F) Overlap of miRNAs determined to be ZSWIM8-sensitive in human fibroblast cell lines with those found to be sensitive MEFs. G) ZSWIM8 sensitivity of miR-335-3p, miR-424, and miR-503 in human fibroblast cell lines. Each point represents the fold-change of normalized expression in ZSWIM8 knockdown cells, relative to the mean of normalized expression in WT samples after normalization to cluster members. (*n* = 3 biological replicates per cell line). Bars represent the mean for each cell line. H) Response of miR-503-5p and miR-424-5p to the knockdown of *PLAGL1*, *LRRC58*, or *ZSWIM8* in BJ-5ta cells (*n* = 2 biological replicates per mRNA); otherwise, as in G.

We next considered the evolutionary conservation of the newly identified TDMD sites. The *Lpar4* and *Atp6v1g1* genes are broadly conserved among vertebrates, but their miR-335 TDMD sites are only present in placental mammals (Figure 1C–D, S7A–C). This conservation is consistent with emergence of the *Mir335* gene after the divergence of placental mammals and marsupials. However, the three TDMD sites exhibit different patterns of conservation within placental mammals. The *Atp6v1g1* site is broadly conserved and is predicted to maintain its TDMD-triggering base pairing pattern in nearly all placental species (Figure 1C, S7B). In a handful of species, such as guinea pigs, a single-nucleotide G-to-A substitution in *Atp6v1g1* across from miRNA nucleotide 15 is predicted to convert a G:U wobble into a canonical base pair and thereby strengthen the pairing to miR-335.

In contrast, the two sites in *Lpar4* have weaker evolutionary conservation (Figure 1D, S7C). In primates, the second site has frequently lost a canonical seed match, but this loss often appears to be compensated by strengthened pairing to the first site, wherein G:U wobbles are replaced with canonical pairing. In contrast, moles lost substantial 3′ pairing in the first site while maintaining extensive complementarity in the second site. Thus, the presence of multiple sites targeting the same miRNA appears to have provided additional opportunity for compensation and for tuning the precise extent of TDMD over mammalian evolution.

Similar to miR-335, the miR-322/503 cluster arose soon after the divergence of placental mammals and marsupials and is conserved in nearly all placental animals. The miR-503 site within *Lrrc58* is also highly conserved throughout placental mammals and is the most conserved segment within the *Lrrc58* 3′ UTR (Figure 2C). Moreover, the interaction between miR-503 and its TDMD trigger site is further supported by covariation, in which a single-nucleotide change at nucleotide 17 of monkey and ape miR-503 is accompanied by a change in the miR-503 binding site that preserves base pairing (Figure S8A). Additionally, the miR-503 of new-world monkeys possesses a change to A at nucleotide 16, which further extends complementarity between miR-503 and *Lrrc58*.

In contrast, the miR-322 trigger site appears to have a more varied evolutionary history. A seed match to miR-322 is present throughout mammalian *Plagl1* sequences, and some 3′ pairing is present in most mammalian species, suggesting that *Plagl1* is typically a miR-322 target and that the ancestral site possessed canonical seed as well as supplementary pairing (Figure S7C). However, the extensive complementarity observed in mouse is present only among members of the *Muridae* family as well as within the *Afrotheria* clade. Within *Muridae*, two sequential changes appear to have led to the acquisition of extensive complementarity: (1) a change of miR-322 nucleotide 21 from the ancestral A to a G in a common ancestor of the *Muroidae* superfamily (including voles, hamsters, and jerboas), which resulted in complementarity to a C nucleotide in *Plagl1*, and (2) a change of the *Plagl1* sequence in a common ancestor of the *Muridae* family, which resulted in complementarity to miR-322 nucleotides 12–14. Thus, the ancestral miR-322 site may have been poised to acquire the ability to direct miR-322 degradation but appears to have acquired it only recently, in an ancestor of mice and rats, as well as in an ancestor of *Afrotheria*. Accordingly, in contrast to miR-335-3p and miR-503, we did not detect ZSWIM8 sensitivity of miR-424, the human ortholog of miR-322, suggesting that *Plagl1* TDMD activity does not extend to humans (Figure 3G). Analogous results were obtained when knocking down newly identified TDMD triggers in BJ-5ta human fibroblasts (a telomerase-immortalized derivative of the BJ cell line). Consistent with the predictions of our evolutionary analysis, we observed upregulation of miR-503 upon *LRRC58* knockdown but did not observe upregulation of miR-424 upon *PLAGL1* knockdown (Figure 3H, S6G–I). (Although ZSWIM8 sensitivity of miR-335-3p is conserved in humans (Figure 3G), we were unable to assess the conservation TDMD by human *ATP6V1G1* and *LPAR4* because miR-335-3p is not expressed in BJ-5ta fibroblasts.)

### *Plagl1-* and *Lrrc58*-directed degradation of miR-322 and miR-503 enhances growth of mice

*Zswim8^-/-^*mice possess growth and developmental defects (Jones et al. 2023; Shi et al. 2023), but how deregulation of miRNA degradation contributes to each defect is unclear. Indeed, some or all of these defects might be attributed to disrupted degradation of non-AGO targets reported for this E3 ligase substrate receptor (Molina-Pelayo et al. 2022; Wang et al. 2023; Ren et al. 2024). To begin to link targeted degradation of specific miRNAs to their downstream consequences, we evaluated the effects of disrupting the TDMD trigger sites for miR-322 and miR-503. Mice with disrupted trigger sites within *Plagl1*, *Lrrc58*, or both triggers were viable and fertile but had reduced body size. At E18.5, *Plagl1* and *Lrrc58* mutant mice were 5.5% and 4.8% smaller by weight compared to wild-type littermates (Figure 4A–B). Furthermore, these effects were additive, as *Plagl1*/*Lrrc58* double-mutant mice were yet smaller — 12% smaller at E18.5 (Figure 4C), which accounts for approximately half of the growth defect observed in *Zswim8^-/-^*mice at this developmental timepoint. The growth defect observed in *Plagl1*, *Lrrc58*, and *Plagl1*/*Lrrc58* double-mutant mice persisted through adulthood; at 8 weeks old, mice expressing mutant *Plagl1* or *Lrrc58* were ∼7–8% smaller, and *Plagl1*/*Lrrc58* double-mutant mice were 14% smaller than control littermates (Figure 7D–F). These results were consistent with a previously reported growth increase in mice with a deletion of the miRNA cluster containing *Mir-322* and *Mir-503* (Jones et al. 2023), but further demonstrate a causal relationship between TDMD and growth.

**Figure 4.**
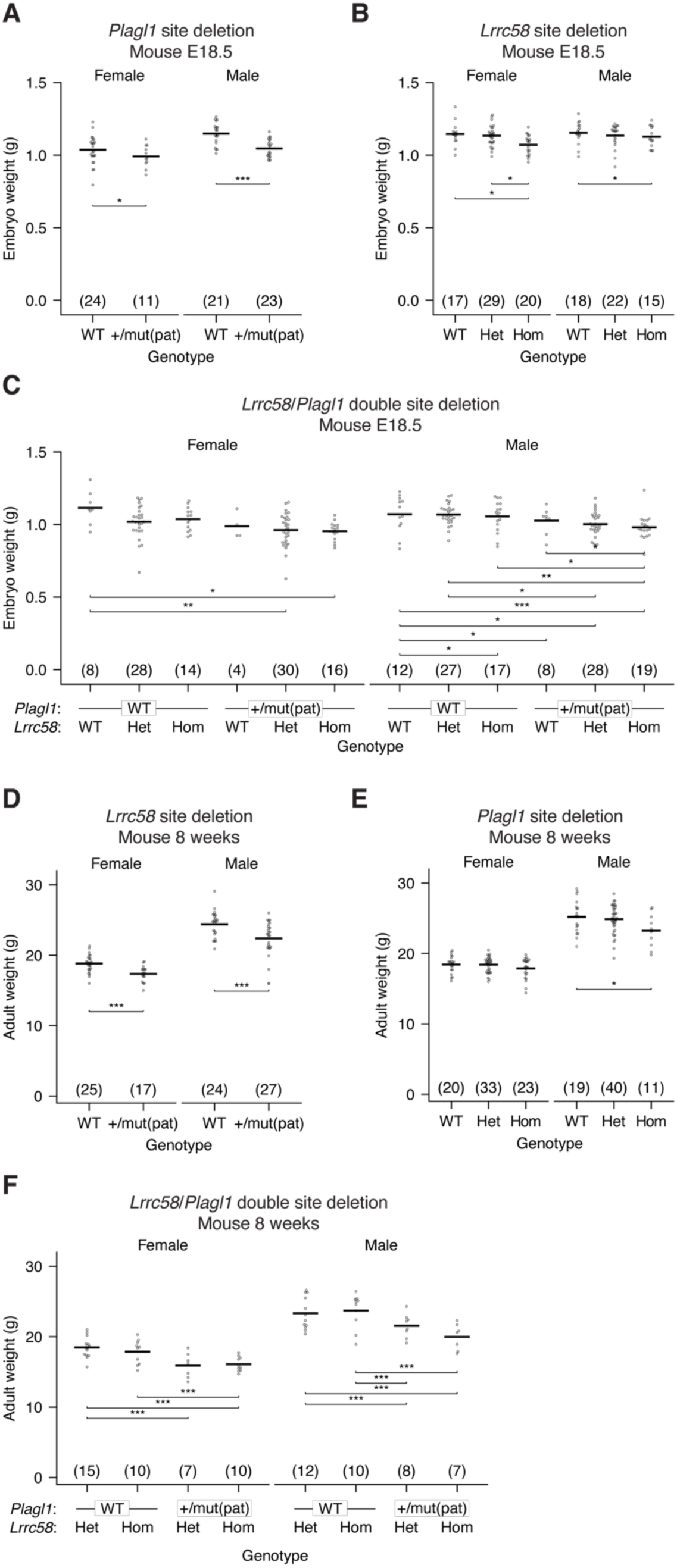
TDMD of miR-322 and miR-503 promotes growth. A) Loss of *Plagl1*-directed miR-322 degradation reduces embryonic growth. Shown are weights of E18.5 mouse embryos with either a wild-type (WT) or a mutant miR-322 trigger site in the expressed *Plagl1* allele. “mut(pat)” indicates a paternally inherited *Plagl1* allele with a mutation in the miR-322 trigger site. The number of animals for each genotype is indicated in parentheses. Statistical testing was performed using a mixed-linear-effects model (* *P* <0.05, ** *P* <0.005, *** *P* <0.0005). B) Loss of *Lrrc58*-directed miR-503 degradation reduces embryonic growth (WT, wild-type; het, heterozygous mutant *Lrrc58*; hom, homozygous mutant *Lrrc58*); otherwise, as in (A). C) Loss of both *Plagl1*-directed miR-322 degradation and *Lrrc58*-directed miR-503 degradation further reduces embryonic growth. Plotted are weights of E18.5 embryos generated from crossing *Lrrc58* het*, Plagl1* WT females with *Lrrc58* het*, Plagl1*^+/mut(pat)^ males; otherwise, as in (A) and (B). D) Loss of *Plagl1*-directed miR-322 degradation reduces the size of adult mice. Shown are weights of 8-week-old mice with either a wild-type (WT) or a mutant miR-322 trigger site at the expressed *Plagl1* allele (+/mut(pat)). Otherwise, as in A. E) Loss of *Lrrc58*-directed miR-503 degradation reduces the size of adult mice. Plotted are weights of 8-week-old mice with either a wild-type, heterozygous mutant, or homozygous mutant miR-503 trigger site within *Lrrc58*; otherwise, as in (B). F) Loss of both *Plagl1*-directed miR-322 degradation and *Lrrc58*-directed miR-503 degradation further reduces the size of adult mice. Plotted are weights of 8-week-old mice generated from crossing *Lrrc58* hom*, Plagl1* WT females with *Lrrc58* het*, Plagl1*^+/mut(pat)^ males; otherwise, as in C.

## Discussion

In this study, we identify five TDMD trigger sites within four trigger transcripts that direct the degradation of three murine miRNAs. These results, some of which have been reported from parallel efforts in other labs (LaVigne et al. 2025; Li et al. 2025), help to triple the number of endogenous TDMD triggers identified in mice. *Plagl1* and *Lrrc58* direct degradation of the related and co-transcribed miR-322 and miR-503, respectively, and *Atp6v1g1* and *Lpar4* collaborate to direct degradation of miR-335-3p. Identifying these trigger transcripts causally linked increased expression of these miRNAs in the *Zswim8* knockout context (i.e., ZSWIM8 sensitivity) to their degradation through the TDMD pathway.

While preparing our manuscript for publication, others reported identification of the trigger sites in *Plagl1*, *Lrrc58*, and *Atp6v1g1,* as well as the *Plagl1*/*Lrrc58*-dependent growth defect (LaVigne et al. 2025; Li et al. 2025). In addition to independently supporting the results of those parallel studies, we identified and validated *in vivo* TDMD activity for two trigger sites within *Lpar4* that collaborate with each other and a site in *Atp6v1g1* to direct the degradation of miR-335-3p, and in doing so, help establish the principles that 1) multiple sites on the same transcript can collaborate to direct the degradation of a miRNA and 2) multiple sites on different transcripts can collaborate to direct the degradation of a miRNA.

The five trigger sites are within the 3′ UTRs of four protein-coding genes with varied functions. *Atp6v1g1* encodes an essential vacuolar ATPase, *Lpar4* encodes a G protein-coupled receptor, *Plagl1* encodes a zinc-finger transcription factor, and *Lrrc58* encodes a recently described E3 ubiquitin ligase substrate adaptor involved in cysteine catabolism (Ramage et al. 2025; Xiao et al. 2025). The trigger sites contained within these four transcripts display a wide range of evolutionary conservation. The *Lrrc58* and *Atp6v1g1* trigger sites emerged early in eutherian evolution and have been broadly conserved across placental mammals. *Lpar4* trigger sites also emerged in early eutherians but are somewhat less conserved, although most placental mammals harbor at least one site, and the *Plagl1* trigger sites appear to have emerged only recently, in the rodent lineage and perhaps independently within *Afrotheria*.

The substitutions that generate a new trigger site provide a mechanism for an mRNA to acquire two independent functions — the ability to code for a protein and the ability to direct miRNA degradation and thereby indirectly influence the expression of hundreds of transcripts. Mutating trigger sites within the 3′ UTR of either *Lrrc58* or *Plagl1* did not significantly change accumulation of the trigger mRNA itself (Figure S5B), indicating a resistance to mediating repression of their host mRNA, also observed for TDMD trigger sites in flies (Kingston et al. 2022) and indicating that the sites do not serve double-duty to repress both the miRNA and the host mRNA.

Although previously identified endogenous trigger RNAs are all highly expressed in the contexts in which they mediate TDMD (Bitetti et al. 2018; Kleaveland et al. 2018; Shi et al. 2020; Kingston et al. 2022), our results indicate that this is not a universal property. *Lpar4* causes detectable TDMD at much lower expression levels than previously reported TDMD triggers (Figure S9). Indeed, *Lpar4* appears to degrade miR-335-3p as much as *Atp6v1g1* in liver, where *Atp6v1g1* is >55-fold more abundant. Further studies will be required to determine whether this unusual potency is due to the particular pairing patterns of the sites in *Lpar4* or whether it is due to other features of this exceptional trigger transcript.

Our computational approach identified all five sites for the three miRNAs of this study within its top-scoring candidates. However, this approach has not been productive for some other ZSWIM8-sensitive miRNAs. This uneven success suggests that not all trigger sites have the features inferred from the initially identified sites used to train our search. Indeed, we observe that *Plagl1*-mediated TDMD is not broadly conserved whereas *Lpar4*-mediated TDMD does not require high expression levels, and recent reports in nematodes and flies indicate that not all trigger–miRNA pairs possess extensive 3′ complementarity (Donnelly et al. 2022; Hiers et al. 2025). For example, in *C. elegans*, a family of miRNAs is rapidly degraded using a mechanism that does not seem to require extensive pairing to the 3′ region of the miRNA (Donnelly et al. 2022; Grimme et al. 2025). Unbiased genetic or biochemical approaches may reveal triggers for more of the ZSWIM8-sensitive mammalian miRNAs.

ZSWIM8 knockout causes lung and heart defects, perinatal lethality, reduced growth, and upregulation of >40 miRNAs in the developing mouse embryo (Jones et al. 2023; Shi et al. 2023). However, when considering that ZSWIM8-directed ubiquitylation might have additional targets beyond AGO, no causal link between TDMD and these phenotypes had been demonstrated. Our work and parallel work by the Mendell lab (LaVigne et al. 2025) causally links the ZSWIM8-dependent growth defect at least partially to target-directed degradation of miR-322 and miR-503. Deletion of the miR-322 and miR-503 trigger sites within *Plagl1* and *Lrrc58* phenocopies 55% of the embryonic growth defect observed in *Zswim8* knockout mice. Characterization of other TDMD trigger mutants will shed light on whether the other reported phenotypes are also caused by dysregulation of miRNA levels and may reveal other physiological roles for regulated miRNA degradation.

miR-322 and miR-503 are members of the extended miR-16 family, which have many known connections to cell cycling and growth. Targets of miR-16 family miRNAs include known cell cycle regulators, such as *BCL2*, MAP kinase pathway members, *CYCLIN D1/2/3*, *CYCLIN E1*, *CDC25A*, and *CDK6* (Cimmino et al. 2005; Linsley et al. 2007; Bonci et al. 2008; Liu et al. 2008; Marasa et al. 2009; Rissland et al. 2011). These miRNAs are also deleted or downregulated in various cancer contexts, where cells are rapidly cycling (Calin et al. 2002, 2005; Bandi et al. 2009; Klein et al. 2010; Liu et al. 2019). Conversely, overexpression of miR-16 family members results in accumulation of cells in G1 (Linsley et al. 2007; Liu et al. 2008). Another validated target of the miR-16 family is *Insulin-like Growth Factor 1 Receptor* (*IGF1R*) (Llobet-Navas et al. 2014). IGF1 is a regulator of fetal growth and development, and defects in IGF1 and IGF1R impair embryonic growth (Baker et al. 1993; Liu et al. 1993; Powell-Braxton et al. 1993). The upregulation of miR-322 and miR-503 and consequent enhanced repression of their target mRNAs in TDMD mutant mice presumably mediates growth restriction through the pathways described above, among others.

Our results showing that newly identified triggers together with their associated miRNAs fall within imprinted gene networks link TDMD to genomic imprinting and do so in a manner consistent with the parental conflict hypothesis of kinship theory. *Plagl1* mRNA, the trigger for miR-322, is maternally imprinted (Kamiya 2000; Piras et al. 2000), suggesting that it acts in the interests of the paternal chromosomes, which it indeed appears to do by two different means: 1) it encodes a transcription factor that acts in a broader imprinted gene network to enhance intrauterine growth through induction of other imprinted genes, including *Igf2*, *Cdkn1c*, *Gnas*, and *H19* (Varrault, 2006; Varrault, 2017; Iglesias-Platas, 2014), and 2) it directs the degradation of miR-322, a growth-inhibiting miRNA, (Figures 2 and 4) (LaVigne et al. 2025). Because the previously described *Plagl1*-knockout mice also lost expression of the miR-322 TDMD site (Varrault et al. 2006), the magnitude of intrauterine growth restriction reported for *Plagl1* knockout likely results from disruption of both these mechanisms. In another imprinted layer in this regulatory network, *Plagl1* is repressed by miRNAs from the *Mirg* miRNA cluster (Whipple et al. 2020), which are imprinted on the paternal allele (Seitz et al. 2003). An orthogonal connection between TDMD and imprinting is the role of the non-imprinted genes *Atp6v1g1* and *Lpar4* in degradation of miR-335, a maternally imprinted miRNA expressed within the intron of *Mest*, which is also imprinted and enhances growth in mice (Hiramuki et al. 2015). Many other ZSWIM8-sensitive miRNAs are imprinted (Bartel 2018; Jones et al. 2023; Shi et al. 2023). Determining how more of these presumed regulatory interactions have been leveraged in parental genome conflict will require further investigation. Nonetheless, all evidence seems to indicate that TDMD, like transcriptional and other posttranscriptional regulatory processes (Cleaton et al. 2014; Peters 2014; Tucci et al. 2019), has been a frequent weapon in the conflict between the parental chromosomes.

## Materials and Methods

### Computational pipeline for identification of candidate triggers

Computational identification and scoring of candidate TDMD trigger sites was performed through multiple searches. In the first search, sites for each miRNA sequence that are annotated as conserved targets in TargetScanMouse Release 8.0 (Agarwal et al. 2015; McGeary et al. 2019) were scored. In a separate search, but using the same scoring scheme, all canonical seed matches within mRNA 3′ UTRs as well as long non-coding sequences annotated in Ensembl GRCm38 version 102 were scored. Candidate genes were filtered for expression in mouse embryonic fibroblasts (MEFs) (Shi et al. 2020). For each site, a 30-nucleotide region upstream of each site was searched for the pairing configuration with the highest extent of complementarity to the 3′ end of the miRNA. Complementarity was scored with the following scheme to identify sites similar to previously identified TDMD triggers: (1) each base pair match (after nucleotide 12) was awarded 1 point; (2) base pairing to the penultimate or third-to-last nucleotides of the miRNA was awarded an additional 0.5 points; (3) G:U wobble pairing was not awarded points, except 0.5 points were awarded if a wobble pair was present at the last position of the miRNA (4) 1 point was deducted for each gap or mismatch; (5) offsets larger than 3 were penalized with a 0.5 point penalty per nucleotide greater than 3. In parallel, the base pairing energy was calculated using the RNAduplex function from the ViennaRNA package for all scored candidate sites (Lorenz et al. 2011).

### Computational analysis of trigger conservation

For each miRNA and TDMD trigger site, the sequence alignment from the mammalian 470-way alignment (https://hgdownload.soe.ucsc.edu/goldenPath/hg38/multiz470way/) was downloaded from UCSC using the Table Browser tool (Karolchik 2004). The mature miRNA sequence for each species was predicted from the genomic sequence. 3′ pairing scores were calculated for each cognate miRNA–trigger site pair using the same computational pipeline as above.

### Tissue culture (cell lines + reagents)

All cells were cultured at 37 °C with 5% CO_2_. MEFs and HEK293T cells were cultured in DMEM supplemented with 10% FBS. HFF-1 cells were cultured in DMEM supplemented with 15% FBS. IMR90 cells were cultured in EMEM supplemented with 10% FBS. BJ cells were cultured in EMEM supplemented with 10% FBS. BJ-5ta cells were cultured in a 4:1 mixture of DMEM and Medium 199 supplemented with 0.01 mg/ml hygromycin B and 10% FBS. MEF-CRISPRi and BJ-5ta-CRISPRi cell lines were generated by transducing MEF or BJ-5ta cell lines with a lentivirus encoding a Zim3-dCas9-P2A-GFP fusion protein and sorting for GFP-positive cells (a gift from Jonathan Weissman) (Replogle et al. 2022). All cells were passaged every 3-6 days to maintain confluency between 10 and 75%, except when cells were grown to contact inhibition immediately prior to harvesting.

### Cas9-mediated site deletion

In initial pilot experiments, we observed substantial clone-to-clone variability in miRNA expression levels. To mitigate the potential influence of this variability on assessment of candidate TDMD triggers, we first generated clonal parental cell lines by transfecting MEFs with a PX458-derived plasmid harboring a non-targeting guide and isolating clonal cell lines from the transfected population. These clonal cell lines were used as the parental cell lines for subsequent experiments. Cas9-mediated site deletions were generated by transfecting a clonal parental MEF cell line with two PX458-derived plasmids harboring guides flanking the candidate TDMD site (Table S3). *Zswim8* knockout clones were generated by transfecting the same parental clonal MEF cell line with a single PX458-derived plasmid harboring a guide targeting *Zswim8*. Control cell lines were generated by transfecting the same parental clonal MEF cell line with a PX458-derived plasmid harboring a non-targeting guide. For all experiments, single GFP-positive cells were sorted two or three days after transfection into 96-well plates containing DMEM supplemented with 20% FBS, 50% conditioned media, and penicillin/streptomycin (Gibco). The characterized cell lines were confirmed to be homozygous for the desired mutations by TOPO cloning of genomic DNA amplicons and Sanger sequencing of multiple clones. All clonal cell lines were grown under contact inhibition for at least 5 days before harvesting.

### CRISPRi knockdown

Lentiviral production and transduction were performed as previously described (Shi et al. 2020). Transfer plasmids encoding single guide RNAs were co-transfected with packaging plasmids in HEK293T cells using the reverse transfection technique. For simultaneous knockdown of *Atp6v1g1* and *Lpar4*, the corresponding guides were delivered with a dual-guide transfer plasmid (Replogle et al. 2022). 1.4 μg of transfer plasmid, 0.94 μg of pCMV-dR8.91 packaging plasmid (a gift from Jonathan Weissman), and 0.47 μg of pMD2.G envelope plasmid (Addgene #12259) were transfected per ∼170,000 cells in a 6-well plate using Lipofectamine 2000 and Opti-MEM. After 72 hours, media was collected and centrifuged at 500 g for 10 minutes to remove debris. For each well of a 6-well plate, 500 μL of virus-containing media was added to CRISPRi-MEFs in culture medium supplemented with polybrene (Santa Cruz) at a final concentration of 1 μg/mL. Plates were centrifuged at 1200 g for 1.5 hours. MEF-CRISPRi cells were grown under 4 μg/mL puromycin selection beginning two days after transduction. Media was refreshed every two days. MEF-CRISPRi cells were grown to contact inhibition for at least 5 days before harvesting. CRISPRi knockdown in BJ-5ta-CRISPRi cells was performed similarly, except puromycin selection was performed at a final concentration of 1 μg/mL and cells were not grown under contact inhibition. Knockdown of target genes was confirmed by reverse transcription–quantitative PCR (RT-qPCR).

### RNA extraction

Total RNA was extracted from cultured cell lines and mouse tissues using TRI Reagent (ThermoFisher). Cultured cell lines were scraped from culture dishes into TRI Reagent. Following euthanasia, mouse tissues were rapidly dissected and flash frozen in liquid N_2_ in Eppendorf tubes. Frozen tissue was transferred to a 50 mL conical tube, 1–2 mL of TRI Reagent was added, and the tissue was homogenized using a TissueRuptor and disposable probes (Qiagen). Following resuspension and/or homogenization, samples were phase separated with 200 μL chloroform (J.T. Baker Analytical) for cultured cell lines or 100 μL 1-bromo-3-chloropropane (Sigma) for tissue. Total RNA was precipitated in isopropanol, washed twice in 75% ethanol, and resuspended in water.

### Small RNA Northern blot

5 μg total RNA was resolved on a denaturing 15% polyacrylamide gel and transferred to Hybond NX or Hybond N+ membranes (Cytiva) using a semi-dry transfer apparatus (Bio-Rad). To crosslink RNA to the membrane, the membrane was incubated in a solution of EDC (N-(3-dimethylaminopropyl)-N′-ethylcarbodiimide; Thermo) diluted in 1-methylimidazole at 60°C for 1 h. Radiolabeled DNA or LNA oligonucleotide probes was incubated overnight in ULTRhyb-Oligo Hybridization Buffer (Invitrogen). Prior to re-probing, hybridized probes were stripped from the membrane by incubation in boiling 0.04% SDS with agitation. A detailed protocol is for small RNA Northern blot analysis is available at http://bartellab.wi.mit.edu/protocols.html. Results were analyzed on a Typhoon phosophimager (Cytiva) and quantified using ImageQuant TL (v8.1.0.0). Northern blot probe sequences and hybridization temperatures are listed in Table S3.

### RT-qPCR

For RT-qPCR experiments, cDNA was prepared from 0.5–1 μg total RNA using the QuantiTect Reverse Transcript Kit (Qiagen) according to manufacturer instructions. qPCR experiments were performed on a Roche LightCycler II instrument using Sybr Green I qPCR master mix. qPCR Primer sequences are listed in Table S3.

### mRNA sequencing and analysis

RNA-seq libraries were prepared from total RNA using the Watchmaker RNA Library Prep Kit (Watchmaker Genomics). Ribosomal RNA depletion was performed using RiboDepletion Oligos (Qiagen) by mixing 14 μL of RNA input, 1 μL FastSelect reagent (Qiagen), and 10 μL Frag & Prime Buffer (Watchmaker Genomics) and incubating under the following conditions: 85°C 10 minutes, 75°C 2 minutes, 70°C 2 minutes, 65°C 2 minutes, 60°C 2 minutes, 55°C 2 minutes, 37°C 2 minutes, 25°C 2 minutes, 4°C. Samples were prepared from the First Strand Synthesis step of the Watchmaker RNA Library Prep Kit (Watchmaker Genomics) onward, according to the manufacturer’s instructions.

Libraries were multiplexed using xGen UDI Primers (IDT) and sequenced on the Illumina NovaSeq platform with paired-end reads. Gene expression quantification was performed using salmon with the --gcBias and --validateMappings options to map to the mouse transcriptome (GRCm38, version 102) (Patro et al. 2017). Only reads mapping to mRNAs or lncRNAs were considered for depth normalization. Differential gene expression analysis was performed using DESeq2 v1.38.3 without use of the lfcShrink function (Love et al. 2014).

### sRNA-seq

Small-RNA sequencing libraries were prepared from 5 μg total RNA. 0.5 fmoles of miR-427-5p (*X. tropicalis*) and 0.5 fmoles of lsy-6-3p (*C. elegans*) were added to each sample as spike-ins. Small RNA species were isolated by excising the gel fragment migrating between 18-nt and 32nt radiolabeled internal standards on a 15% polyacrylamide urea gel. Size selected RNA was eluted from the gel, precipitated in ethanol, and ligated to a preadenylated 3′ adapter (AppNNNNTCGTATGCCGTCTTCTGCTTGddC) using T4 RNA Ligase 2 KQ mutant (NEB) in a reaction supplemented with 10% polyethylene glycol (PEG 8000, NEB). The 3′ adapter had 4 random-sequence positions at its 5′ end to reduce ligation bias. Ligated small RNAs were isolated on a 10% polyacrylamide urea gel, precipitated in ethanol, and ligated to a 5′ adapter (GUUCAGAGUUCUACAGUCCGACGAUCNNNN) using T4 RNA Ligase I (NEB) in a reaction supplemented with 10% PEG. The 5′ adapter had 4 random-sequence positions at its 3′ end to reduce ligation bias. Ligated small RNAs were isolated on an 8% polyacrylamide urea gel, precipitated in ethanol, and reverse transcribed with SuperScript III (Invitrogen). Resulting cDNA was amplified using KAPA HiFi DNA polymerase (Kapa Biosystems). Amplified DNA was purified on a 90% formamide, 8% acrylamide gel and submitted for sequencing on the Illumina HiSeq or NovaSeq platform. A step-by-step protocol for constructing libraries for small-RNA sequencing is available at http://bartellab.wi.mit.edu/protocols.html.

Adaptor sequences were trimmed from reads using cutadapt (Martin 2011). Reads were filtered for quality using fastq_quality_filter (FastX Toolkit; http://hannonlab.cshl.edu/fastx_toolkit/) with the parameters “–q 30 –p 100.”

To assign processed sequencing reads to miRNAs, the first 19nt of each read was matched to a dictionary of miRNA sequences downloaded from TargetScan Release 8.0 (Agarwal et al. 2015; McGeary et al. 2019), requiring no mismatches between the read and the miRNA dictionary. Reads mapping to the spike-in miRNAs and markers were removed for further analysis. Differential expression analysis was performed using DESeq2 v1.38.3 without use of the lfcShrink function (Love et al. 2014).

### miRNA targeting

miRNA targeting analysis was performed as described in Stefano et al, 2025 (Stefano et al. 2025). Briefly, miRNA target predictions were downloaded from TargetScan Release 8.0 (Agarwal et al. 2015; McGeary et al. 2019), and repression of predicted miRNA targets was analyzed in differential expression data. Genes expressed at fewer than 10 TPMs across samples were excluded. Two sets of predicted targets were analyzed: all predicted targets and top predicted targets (10% of targets with the lowest cumulative weighted context++ scores) (Agarwal et al. 2015). Each set of targets was compared to a control group of genes not predicted to be targets of the miRNA family under consideration. The nontarget cohort was selected by sampling transcripts at a one-to-one ratio with targets, matching the distribution of 3′ UTR lengths between the target and nontarget cohorts. The distribution of log2 fold changes (DESeq2 output) in samples relative to *Plagl1* WT, *Lrrc58* het samples was compared to that of the nontarget cohort. Statistical significance was assessed using a Mann–Whitney U test. The degree of repression is represented by subtracting the median log2 fold change of the target set from that of its corresponding nontarget set. The mean degree of repression and median *P* value across 21 iterations of the above analysis are reported. Figure S5C displays a representative cumulative distribution function (derived from the iteration that generated the median *P* value for the all targets set), with only the nontarget set corresponding to the all targets set shown for simplicity.

### Mouse husbandry

Mice were housed at the Whitehead Institute for Biomedical research in accordance with protocols approved by the Massachusetts Institute of Technology Committee on Animal Care. Mice were housed in a 12-hour light/dark cycle (light from 7:00–19:00) with free access to food and water. Euthanasia of adults was performed by CO_2_ inhalation; euthanasia of embryos was performed by rapid decapitation over ice.

### Generation of mutant mice

Mutant mice were generated by the Whitehead Institute Genetically Engineered Models core. Mice with mutations in *Atp6v1g1, Lpar4, Plagl1,* and *Lrrc58* 3′ UTRs were generated by injecting or electroporating C57BL/6J embryos with Cas9 protein complexed with a sgRNA designed to cut within the regions of the 3′ UTR predicted to engage in TDMD (see Table S3). For *Plagl1* and *Lrrc58*, 1-cell embryos were electroporated with Cas9, sgRNAs, and HDR donor oligos. For *Lpar4*, Cas9, sgRNAs, and HDR donor were injected into 1-cell embryos. For *Atp6v1g1*, Cas9 and sgRNAs were injected in 1 blastomere of 2-cell embryos. F_0_ mice containing resulting deletions and mutations were bred to C57BL/6J mice and then backcrossed for at least 2 generations to obtain the desired heterozygous mice used to generate embryos and adult mice used in this study. Mutant lines were maintained by breeding to C57BL/6J or heterozygotes. No substantial phenotypic differences were observed between mice bearing different mutant alleles in *Plagl1* or *Lrrc58*, so mutant alleles were used interchangeably in this study.

### Genotyping

Initial genotyping of mutant mice generated for this study was performed by extracting genomic DNA using the HotSHOT method (Truett et al. 2000), amplifying regions of interest by PCR using the primers listed in Table S3, and analyzing genomic sequences by nanopore sequencing. *Atp6v1g1* mutant mice were genotyped by gel electrophoresis of PCR amplicons containing the TDMD site for all subsequent studies. For other mutant mice, automated genotyping was performed by Transnetyx (Cordova, TN).

### Timed mating and tissue collection

Pregnant females were euthanized by CO^2^ inhalation at embryonic day 18.5 (E18.5; 18 days after inspection of a vaginal plug). Embryos were rapidly dissected over ice and weighed after gently blotting dry. Embryos were decapitated and tissues dissected in ice-cold PBS. Tissues were flash frozen and carried forward for RNA extraction as described.

### Mouse weight analysis

Mutant mice and their littermates were weighed at E18.5 or 8 weeks of age. Statistical testing for mouse weights was performed separately for each sex by using a linear-mixed-effects model, with the litter as a random effect and genotype as a linear effect. Reported *P* values are the pairwise comparisons between genotypes with Tukey correction.

## Acknowledgements

We thank Lianne Blodgett, Elena Slobodyanyuk, Michelle Frank, Peter Wang, Charlie Shi, Elena Kingston, and other current and former members of the Bartel lab for helpful discussions. We thank Jonathan Weissman for sharing CRISPRi-related plasmids, Mohamed El-Brolosy for advice, Kristin Bahleda for assistance for animal husbandry, and Lianne Blodgett and George Bell for sharing computational scripts. We thank the Whitehead Institute Genome Technology Core for sequencing, the Whitehead Institute Flow Cytometry Core for support with cell sorting, the Whitehead Bioinformatics and Research Computing core for advice about computational analyses, and the Whitehead Institute Genetically Engineered Models Center for generation of mutant mouse lines. The project described was supported by NIH grants R35GM118135 (D.P.B.), T32GM007753 (L.E.E), T32GM144273 (L.E.E), and T32GM136540 (L.E.E) from the National Institute of General Medical Sciences, and F30HL175923 (L.E.E.) from the National Heart, Lung, and Blood Institute. The content is solely the responsibility of the authors and does not necessarily represent the official views of the National Institute of General Medical Sciences, the National Heart, Lung, and Blood Institute, or the National Institutes of Health. D.H.L. was a Howard Hughes Medical Institute Fellow of the Damon Runyon Cancer Research Foundation (DRG-2345-18). D.P.B is an investigator of the Howard Hughes Medical Institute.

## Author Contributions

D.H.L. and D.P.B. conceived the project. D.H.L. and L.E.E. performed experiments and analyzed data. E.K. wrote the computational pipeline for identification of trigger candidates. D.H.L., L.E.E., and D.P.B. wrote the manuscript, and all authors edited the manuscript.

---

**Figure S1.**
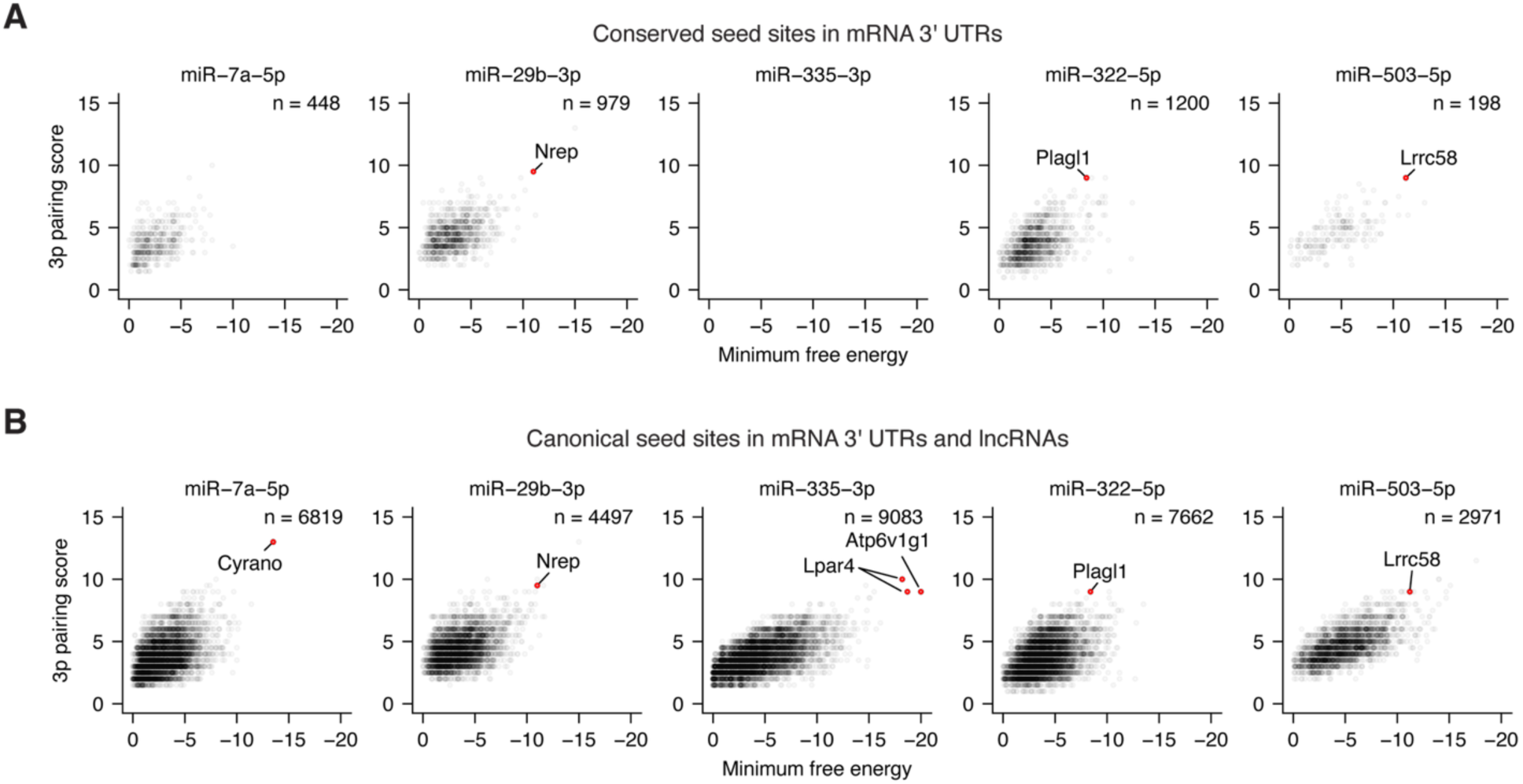
A computational pipeline predicts TDMD trigger sites for miR-335-3p, miR-322-5p, and miR-503-5p. A) Results of a version of the trigger-site prediction pipeline that scores conserved sites. For each conserved seed site (8mer, 7mer-A1, 7mer-m8) in 3′ UTRs, the 3′-pairing score is plotted as a function of predicted 3′-pairing energy. Validated and newly identified TDMD trigger sites are labeled and highlighted in red. miR-335-3p was not analyzed using this pipeline because it had no conserved sites annotated by TargetScan, as it was considered a passenger strand when conserved sites were annotated by TargetScan, and TargetScan does not predict conserved sites of passenger strands. B) Results of a version of the trigger-site prediction pipeline that also scores all non-conserved seed sites in 3′ UTRs and all seed sites in lncRNAs; otherwise, as in A.

**Figure S2.**
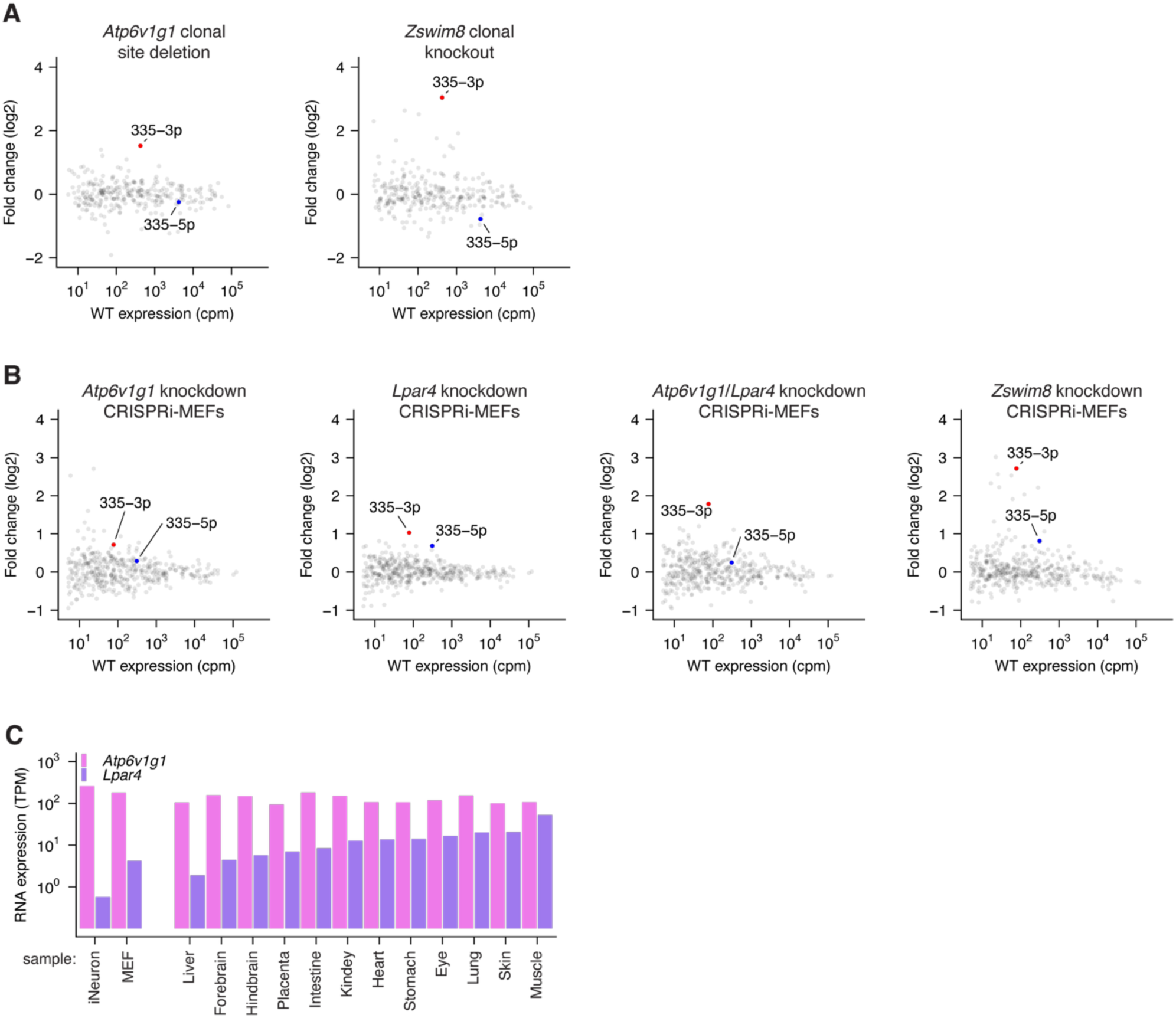
sRNA-seq from cell lines validates *Atp6v1g1* and *Lpar4* as endogenous TDMD triggers. A) sRNA-seq results showing the changes in miRNA levels observed upon disruption of either the miR-335-3p trigger site within *Atp6v1g1* or the *Zswim8* gene. Results from 3–4 clonal lines per genotype were analyzed using DESeq2. The point for miR-335-3p is in red, and the point for the coproduced miR-335-5p strand is in blue. B) sRNA-seq results showing changes in miRNA levels observed when CRISPRi-MEFs were transduced with guides targeting either *Atp6v1g1*, *Lpar4*, both *Atp6v1g1* and *Lpar4*, or *Zswim8*, compared to cells targeted with non-targeting guides. Results from two biological replicates were analyzed using DESeq2; otherwise, as in (A). C) Relative expression of *Atp6v1g1* and *Lpar4* mRNAs, as quantified by RNA-seq in mouse cell lines and embryonic tissues (Shi et al. 2023).

**Figure S3.**
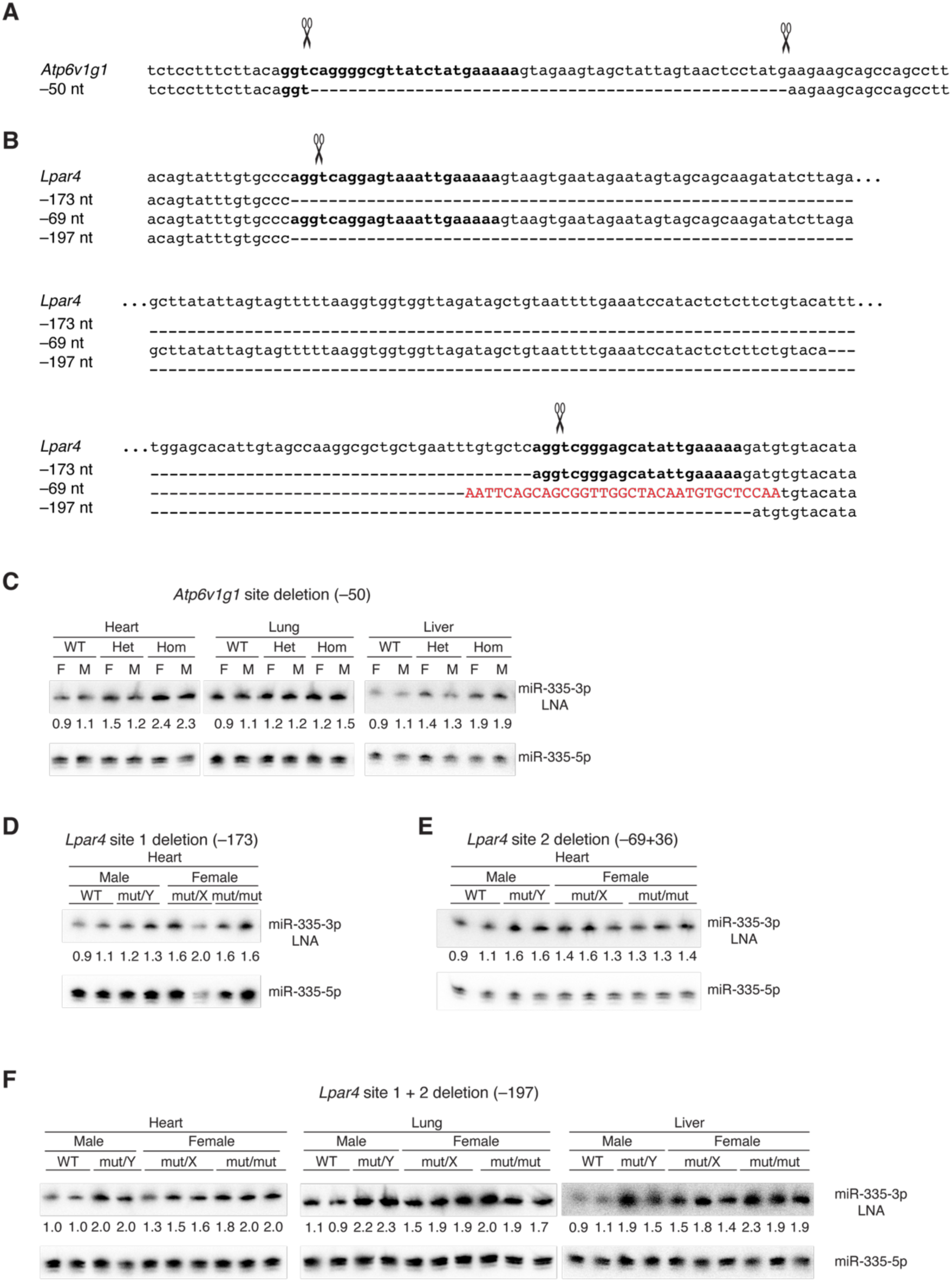
Validation of *Atp6v1g1* and *Lpar4* as TDMD triggers for miR-335 in mice. A) Wild-type and mutant sequences of mouse *Atp6v1g1*. For each gene, the wild-type sequence is shown above the mutant sequence. TDMD sites are in bold; gaps are indicated by dashes; substitutions are in red, and predicted Cas9 cleavage sites are indicated by scissors above the wild type sequences. B) Wild-type and mutant sequences for *Lpar4*; otherwise, as in A. C) Validation of *Atp6v1g1* as a TDMD trigger for miR-355-3p in mice. Shown is a northern blot resolving total RNA of heart, lung, and liver of E18.5 mice, probed for miR-335-3p. Mice were either wild-type (WT), heterozygous, or homozygous for a deletion of the TDMD trigger site in *Atp6v1g1*. Numbers below the lanes indicate the relative fold change, which was obtained by normalizing first to the value of the miR-335-5p loading control and then to the average of the WT samples. D–E) Validation of *Lpar4* as a TDMD trigger for miR-355-3p in mice. Shown are northern blots of RNA from E18.5 heart of *Lpar4* mutant mice in which either TDMD site 1 (*Lpar4*^-173^) or TDMD site 2 (*Lpar4*^-69+36^) has been deleted (panels D and E, respectively); otherwise, as in panel C. F) Effect of deleting both trigger sites within *Lpar4*. Shown are northern blots of RNA from E18.5 heart, lung, and liver for *Lpar4* mutant mice in which both TDMD sites have been deleted (*Lpar4*^-197^). Otherwise, as in panel C.

**Figure S4.**
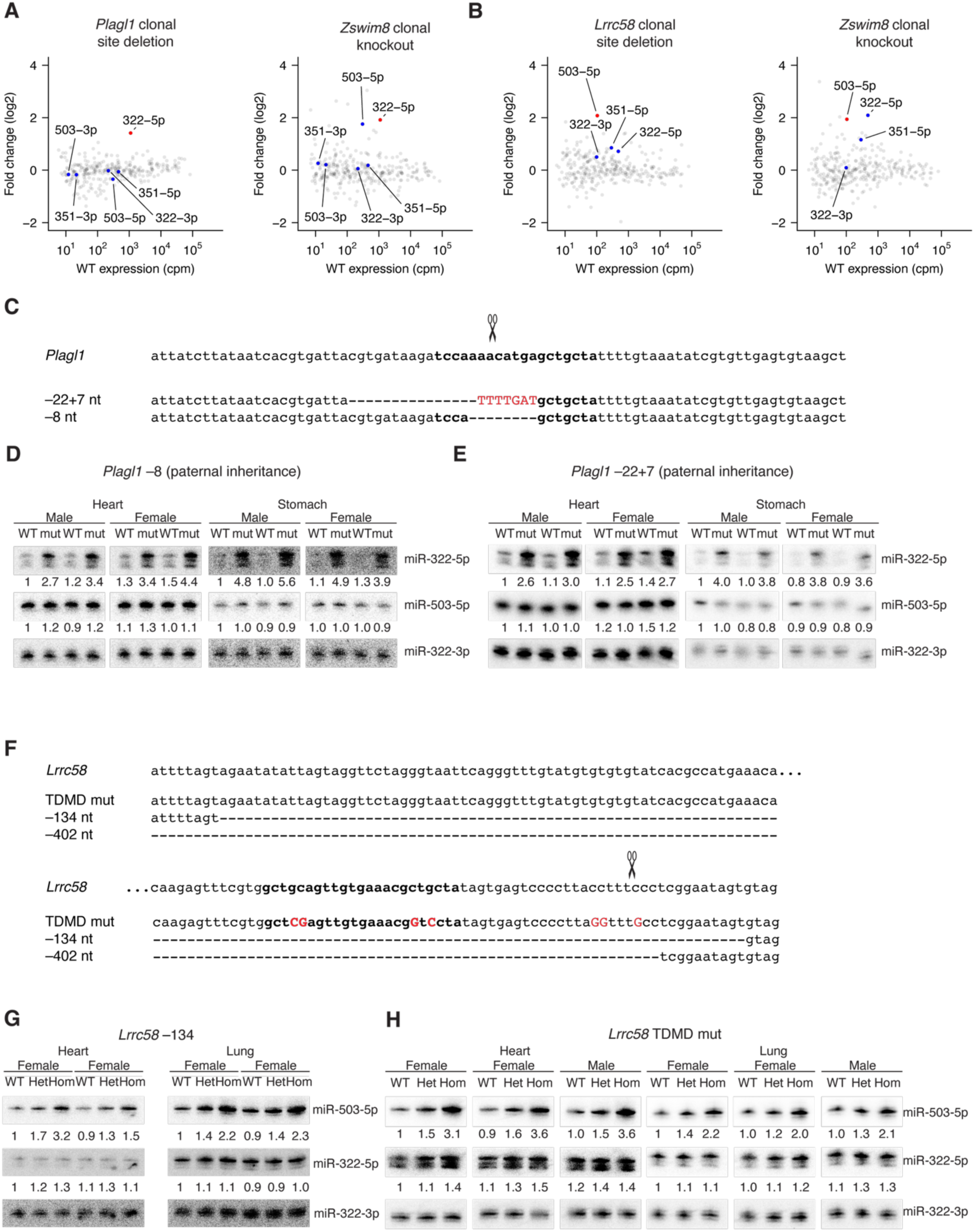
Validation of *Plagl1* and *Lrrc58* as TDMD triggers for miR-322 and miR-503, respectively. A) sRNA-seq results showing the changes in miRNA levels observed upon disruption of either the *Zswim8* gene or the miR-322 TDMD trigger site within *Plagl1*. The point for miR-322-5p is in red, and points for coproduced miR-322-3p, miR-503-5p, miR-503-3p, miR-351-5p, and miR-351-3p are in blue. Results from 3–4 clonal lines per genotype were analyzed using DESeq2. B) sRNA-seq results showing the changes in miRNA levels observed upon disruption of either the *Zswim8* gene or the miR-503 TDMD trigger site in *Lrrc58*. The point for miR-503-5p is in red, and points for coproduced miR-322-5p, miR-322-3p, miR-503-3p, miR-351-5p, and miR-351-3p are in blue. Results from 2–3 clonal lines per genotype were analyzed using DESeq2. C) Wild-type and mutant sequences of mouse *Plagl1*; otherwise, as in Figure S3A. D–E) Validation of *Plagl1* as a TDMD trigger for miR-322-5p in mice. Shown are northern blots of RNA from heart and lung of E18.5 mice with different alleles of trigger-site mutants in *Plagl1* (panels D and E, respectively). Analyses of sex- and litter-matched samples are shown for each of the indicated genotypes. Numbers below the lanes indicate the relative fold change, which was obtained by normalizing first to the value of the loading control (miR-322-3p) and then to the average of the WT samples. F) Wild-type and mutant sequences of mouse *Lrrc58*; otherwise, as in Figure S3A. G–H) Validation of *Lrrc58* as a TDMD trigger for miR-503-5p in mice. Shown are northern blots of RNA from heart and lung of E18.5 mice with different alleles of trigger-site mutants in *Lrrc58* (panels G and H, respectively). Otherwise, as in (D–E).

**Figure S5.**
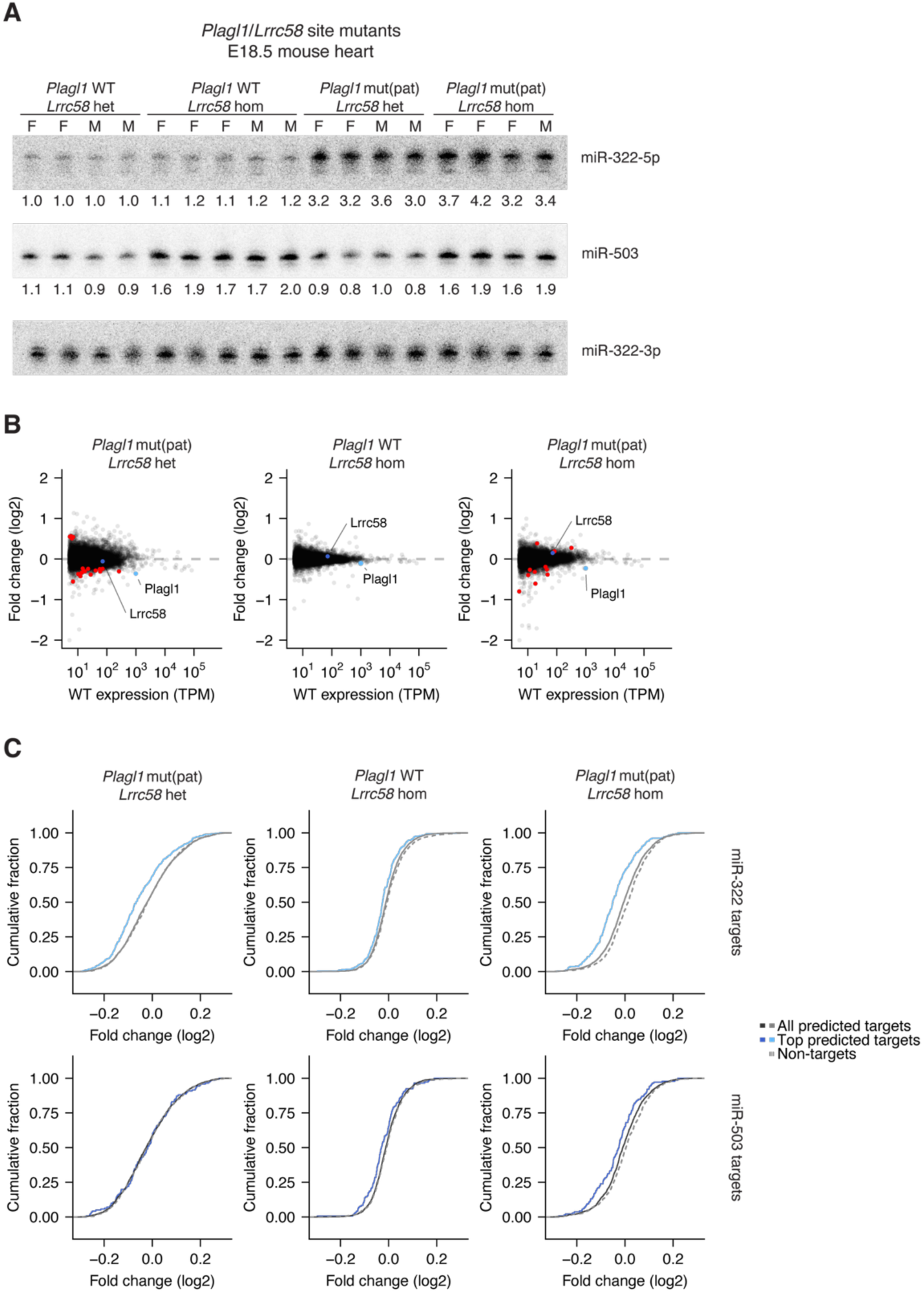
*Plagl1* and *Lrrc58* TDMD trigger sites influence miRNA and mRNA expression in mice. A) Effects of mutating *Plagl1* and *Lrrc58* trigger sites in mice. Shown are northern blots of RNA from heart of E18.5 mice with the indicated trigger site mutants in *Plagl1* and *Lrrc58*. Genotype and sex are indicated above the blots (WT, wild-type; het, heterozygous mutant; hom, homozygous mutant; mut(pat), heterozygous with paternally inherited mutant allele; F, female; M, male). B) RNA-seq results showing changes in mRNA levels observed in mutants of the TDMD trigger sites in *Plagl1* (*Plagl1* mut(pat), *Lrrc58* het), *Lrrc58* (*Plagl1* WT, *Lrrc58* hom), and both *Plagl1* and *Lrrc58* (*Plagl1* mut(pat) *Lrrc58* hom), compared to *Plagl1* WT, *Lrrc58* het littermates. Results from 24 animals (5–7 per genotype) were analyzed using DESeq2. C) The influence of trigger sites within *Plagl1* and *Lrrc58* on levels of miR-503-5p and miR-322-5p predicted targets in vivo. Plotted are cumulative distributions of mRNA fold changes observed in the mutant E18.5 mouse heart relative to wild-type for all predicted targets and top 10% of predicted targets, as determined by TargetScan (Agarwal et al. 2015), and their corresponding control cohorts, matched for 3′ UTR length. For simplicity, only the control cohort corresponding to all predicted targets is displayed (dashed line). The selection of control cohorts was repeated 21 times, and the cohort with the median *P* value (Mann–Whitney U test) is shown here, with this *P* value and the distributions of differences in the median fold changes reported in Figure 2I.

**Figure S6.**
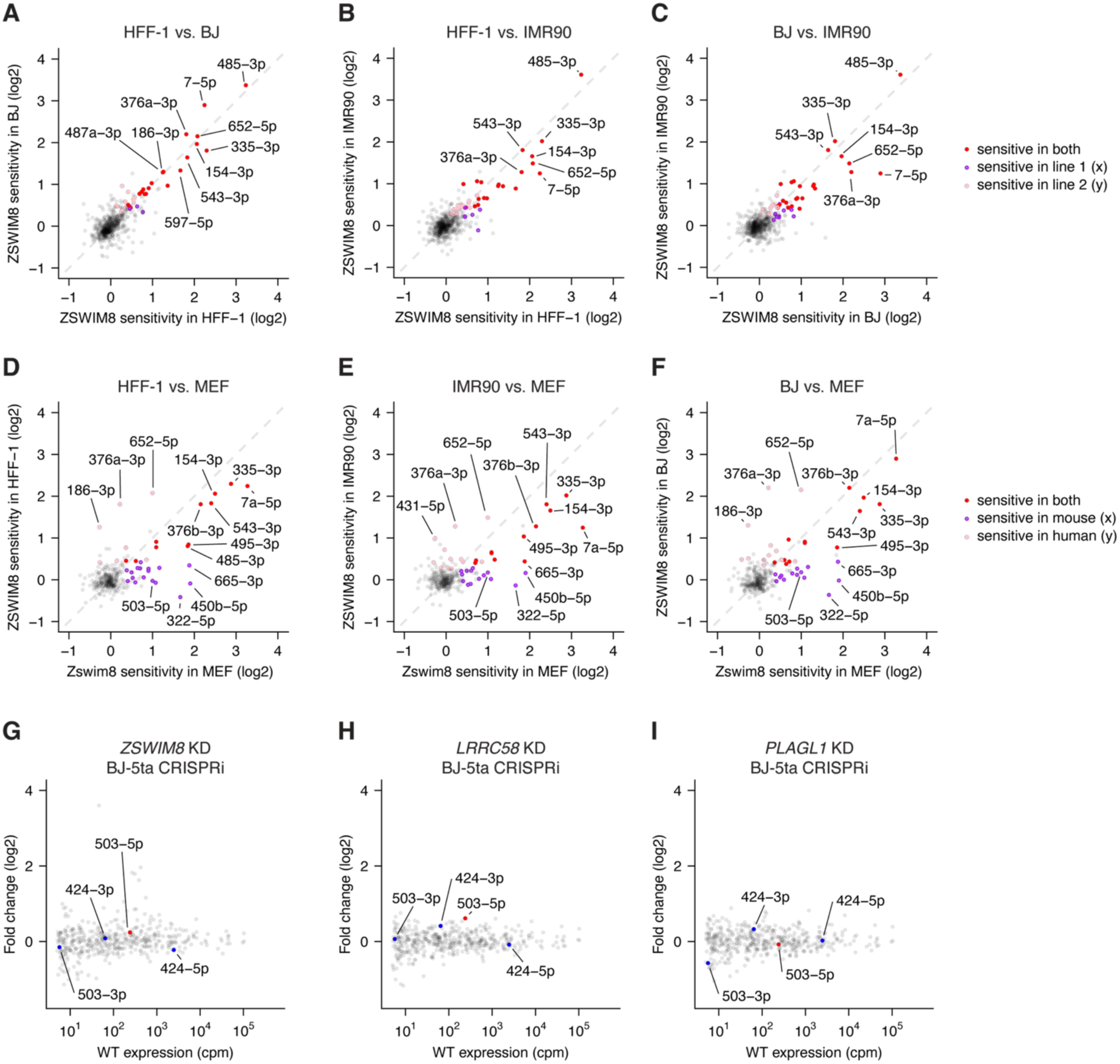
ZSWIM8-sensitve miRNAs are abundant in human fibroblasts. A–C) Abundant ZSWIM8-sensitive miRNAs in three human fibroblast lines. Shown are pairwise comparisons of miRNA fold-changes observed in three different human fibroblast cell lines upon ZSWIM8 knockout, as calculated by DESeq2 (Table S1). Points for miRNAs with statistically significant increases in both cell lines are red (Wang and Bartel 2023). Points for miRNAs with statistically significant increases in only one cell line are purple or pink. D–F) Comparison of ZSWIM8 sensitivities observed between human and mouse. For each miRNA conserved from human to mouse, the fold-change observed in human fibroblast lines upon ZSWIM8 knockout is plotted as a function of the change reported in mouse embryonic fibroblasts (Shi et al. 2023). Colors are as in A. G–H) Conservation of TDMD trigger activity to human cells. Shown are sRNA-seq results plotting miRNA fold changes observed in BJ-5ta CRISPRi human fibroblasts upon knockdown (KD) of either *ZSWIM8* (panel G), *LRRC58* (panel H), or *PLAGL1* (panel I). Results from two biological replicates were analyzed by DESeq2; otherwise, as in Figure S2B.

**Figure S7.**
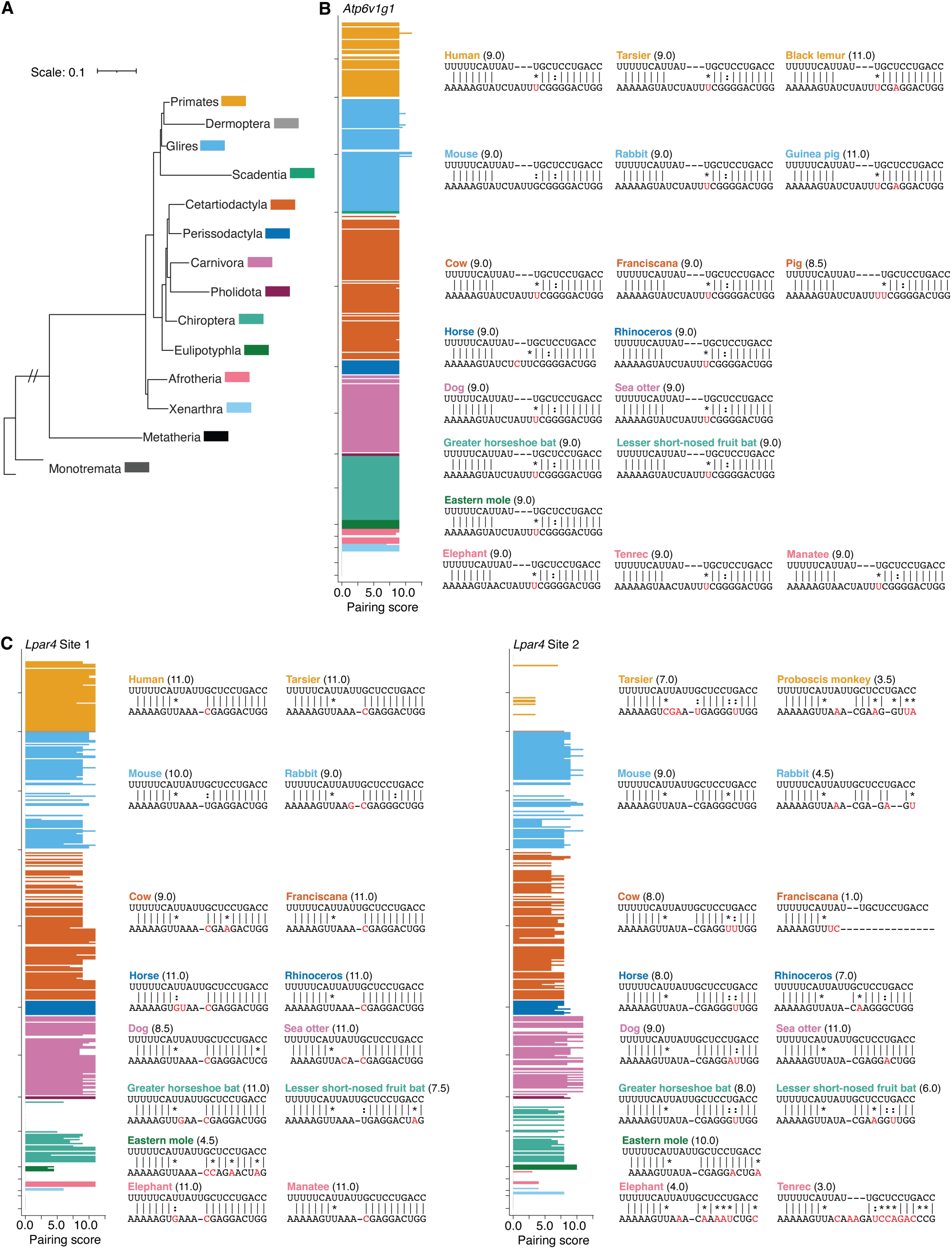
TDMD sites in *Atp6v1g1* and *Lpar4* are evolutionarily conserved. A) Simplified phylogenetic tree of mammalian lineages, based on a 470-species whole-genome alignment (https://hgdownload.soe.ucsc.edu/goldenPath/hg38/multiz470way/) (Letunic and Bork 2024). B) Evolutionary conservation of extensive 3′ complementarity within the miR-335-3p TDMD trigger site in *Atp6v1g1*, across a 470-species mammalian alignment. Each row in the plot corresponds to the 3′ pairing score of our computational pipeline for a species in the 470-way alignment, colored according to clade, as indicated in (A). Missing rows indicate species lacking an orthologous seed match to miR-335-3p. Representative pairing diagrams are shown on the right, with positions that differ from the mouse sequences colored in red. Vertical lines indicate W–C–F pairing; a colon indicates G:U wobble pairing; an asterisk indicates a mismatch. C) As in (B) but for the 2 miR-335-3p trigger sites within *Lpar4*.

**Figure S8.**
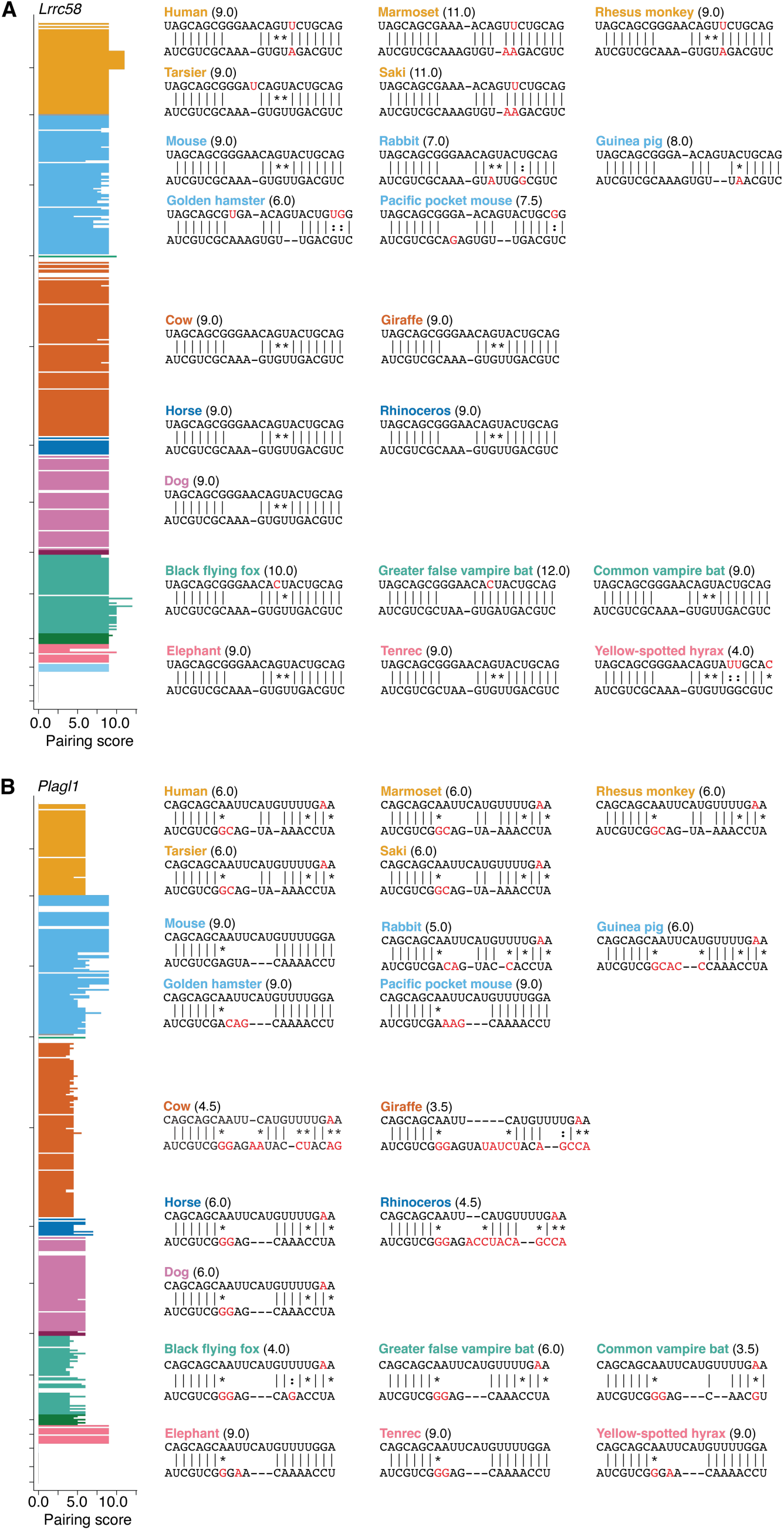
The TDMD sites in *Lrrc58* is more conserved than that in *Plagl1*. A) Evolutionary conservation of extensive 3′ complementarity within the miR-503 TDMD trigger site in *Lrrc58,* across a 470-species mammalian alignment (https://hgdownload.soe.ucsc.edu/goldenPath/hg38/multiz470way/). Missing rows indicate species lacking an orthologous seed match to miR-503. Otherwise, as in Figure S7A. B) As in (A) but for the miR-322/424 site within *Plagl1*.

**Figure S9.**
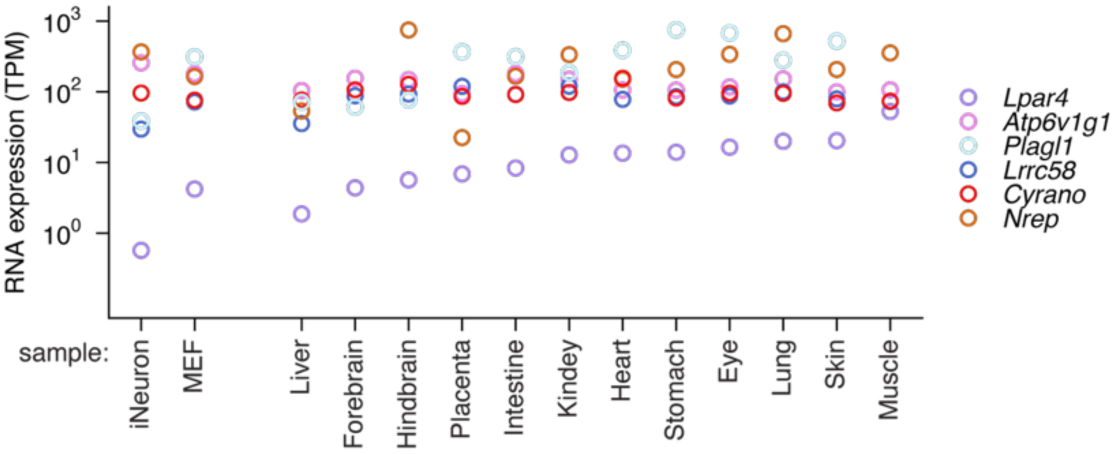
Expression of mammalian TDMD triggers can vary. Relative expression of the six known endogenous mammalian TDMD trigger transcripts, as quantified by RNA-seq in mouse cell lines and embryonic tissues (Shi et al. 2023).

